# Genome-scale CRISPR screening reveals that C3aR signaling is critical for rapid capture of fungi by macrophages

**DOI:** 10.1101/2021.12.30.474615

**Authors:** A. Cohen, E.E. Jeng, M. Voorhies, J. Symington, N. Ali, M.C. Bassik, A. Sil

**Affiliations:** University of California San Francisco, CA, Department of Microbiology and Immunology; Stanford University, Palo Alto, CA, Department of Genetics; AbbVie Inc, San Francisco, CA, Oncology Early Development

## Abstract

The fungal pathogen *Histoplasma capsulatum* (*Hc*) invades, replicates within, and destroys macrophages. To interrogate the molecular mechanisms underlying this interaction, we conducted a host-directed CRISPR-Cas9 screen and identified 361 genes that modify macrophage susceptibility to *Hc* infection, greatly expanding our understanding of host gene networks targeted by *Hc*. We identified pathways that have not been previously implicated in *Hc* interaction with macrophages, including the ragulator complex (involved in nutrient stress sensing), glycosylation enzymes, protein degradation machinery, mitochondrial respiration genes, solute transporters, and the ER membrane complex (EMC). The highest scoring protective hits included the complement C3a receptor (C3aR), a G-protein coupled receptor (GPCR) that recognizes the complement fragment C3a. Although it is known that the complement system reacts with the fungal surface, leading to opsonization and release of small peptide fragments such as C3a, a role for C3aR in macrophage susceptibility to fungi has not been elucidated. We demonstrated that whereas C3aR is dispensable for macrophage phagocytosis of bacteria and latex beads, it is critical for optimal macrophage capture of pathogenic fungi, including *Hc*, the ubiquitous fungal pathogen *Candida albicans*, and the causative agent of Valley Fever *Coccidioides posadasii*. We showed that C3aR localizes to the early phagosome during *H. capsulatum* infection where it coordinates the formation of actin-rich membrane protrusions that promote *Hc* capture. We also showed that the EMC promotes surface expression of C3aR, likely explaining its identification in our screen. Taken together, our results provide new insight into host processes that affect *Hc*-macrophage interactions and uncover a novel and specific role for C3aR in macrophage recognition of fungi.

## Introduction

*Histoplasma capsulatum* (*Hc*) is a fungal intracellular pathogen of macrophages. Infection with *Hc* occurs when soil containing *Hc* spores or hyphal fragments is aerosolized and fungal particles are inhaled by a mammalian host (1). In the lung, *Hc* invades alveolar macrophages (2, 3), replicates to high intracellular levels and induces macrophage lysis (4, 5). Though many of the molecular mechanisms underpinning *Hc* pathogenesis are unknown, a number of *Hc* genes that promote immune evasion and virulence have been identified (6–10).

The initial step in *Hc*-macrophage interactions is phagocytosis. In general, macrophage-expressed pattern-recognition-receptors can directly bind common fungal cell-wall components (11) such as the cell-wall sugar *β*-glucan, which is recognized by the receptor Dectin-1 (12, 13). Engagement of phagocytosis receptors, such as Dectin-1, triggers a complex cascade of intracellular signaling events, involving small GTPase activation, membrane phospholipid remodeling, and actin cytoskeleton polymerization that allow the plasma membrane to deform and encircle the targeted particle (14, 15). Following phagocytosis, the particle is enclosed within a membrane structure termed the phagosome. Macrophage phagocytosis of *Hc*, unlike that of other fungi, is not dependent on *β*-glucan recognition by Dectin-1 (16). *Hc* can prevent such recognition by shielding cell-wall *β*-glucan with a layer of *α*-glucan (17) or by secreting glucanases to prune *β*-glucans (18). Instead, *Hc* recognition and phagocytosis is directly mediated by *β*2 integrin receptors (16, 19) formed through dimerization of CD18 (*Itgb2*) with various *α* subunits (20), such as CD11b, a subunit of complement receptor 3 (CR3).

Innate immune recognition of pathogens is supported by the complement system (21). Dozens of complement factors are secreted into biological fluids such as serum and bronchoalveolar fluid (22, 23), where they react with foreign particles and facilitate their destruction and recognition by innate immune cells (24). The complement cascade can be triggered by three main pathways: the antibody-dependent classical pathway; the lectin pathway, through recognition of microbial sugars; and the non-specific alternative pathway, all of which culminate in the cleavage and activation of C3 (21). Following C3 activation, C3b is covalently attached to the microbial surface, and is recognized by complement receptors (CRs) expressed on immune cells, which mediate phagocytosis (25). C3 cleavage also releases the small peptide fragment, C3a, which is recognized by the complement 3a receptor (C3aR), a G-protein coupled receptor (GPCR) which is expressed on innate immune cells (26). C3a acts as a chemoattractant for innate immune cells such as macrophages (27). C3aR can also modulate the production of cytokines in response to inflammatory stimuli (28), and has been implicated in the pathogenesis of diseases such as sepsis and allergic inflammation (29). C5 is activated downstream of C3, leading to the release of C5a, which is also a potent chemoattractant and inflammatory modifier through its interaction with its receptor, C5aR (29). While serum is a major source of complement, innate immune cells such as macrophages can also secrete complement components (30–32).

Ubiquitous opportunistic fungal pathogens, including *Candida albicans*, as well as endemic fungal pathogens such as *Coccidioides immitis* (33) and *Hc* (34), are strong activators of multiple serum complement pathways (35). Serum enhances the phagocytosis of opportunistic fungal pathogens, and the role of C3b/ inactivated C3b (iC3b) opsonization in promoting uptake of fungi due to recognition by complement receptors is well-studied (35–38). In addition, complement plays an important role in host defense against opportunistic fungi, including *Candida albicans* (39) and *Cryptococcus neoformans* (40). Zymosan, a cell-wall preparation of *Saccharomyces cerevisiae*, is well-established as a model for complement activation (41). Additionally, CR3 can recognize glucans and is thought to promote complement-independent recognition of *Hc* (16, 19, 25). C5a-C5aR signaling can also promote neutrophil migration towards and phagocytosis of *Cryptococcus neoformans* (42) and promote monocyte cytokine production in response to *Candida albicans* infection (43). However, the role of complement in innate immune recognition of *Hc* and of C3a-C3aR signaling in macrophage interaction with fungi has not been investigated.

To characterize host genes that underlie macrophage susceptibility to infection with *Hc*, we took advantage of a powerful pooled host-side screening platform (44) that has been successfully employed to identify host targets of intracellular pathogens (45, 46) and microbial toxins (47). We screened a CRISPR-Cas9 knockout library in macrophage-like cells challenged with *Hc*, and identified genes required for macrophage susceptibility to *Hc*-mediated lysis. We identified a number of host pathways that affected macrophage susceptibility to *Hc* infection, and focused our studies on molecules that influence *Hc* phagocytosis. This led to the discovery that C3aR and GPCR signaling plays an important role in promoting serum-dependent phagocytosis of *Hc* and other fungi. In addition, our screen identified the ER membrane (EMC) complex subunit Emc1, which we discovered is critical for surface expression of C3aR but not integrin receptors. This finding suggests a role for the EMC, which facilitates folding of transmembrane helices in the ER, in the biogenesis of GPCRs in innate immune cells. Overall our findings shed light on molecular mechanisms underlying innate immune recognition of fungi, and uncover new host pathways that may be targeted by *Hc* to promote virulence.

## Results

### A large-scale pooled CRISPR screen in J774A.1 macrophage-like cells identified genes required for macrophage susceptibility to infection with *H. capsulatum*

To identify genes that affect macrophage sensitivity to parisitization by *Hc*, we conducted pooled CRISPR-Cas9 knockout screens in the J774A.1 mouse macrophage-like cell-line (Fig. 1). This cell line has been widely used to model macrophage interactions with pathogenic microbes, including *Hc* (7, 9). We demonstrated that *Hc* can induce lysis of J774A.1 cells in a manner dependent on the secreted effector Cbp1 (Fig. S1A), which is consistent with studies in primary macrophages (7, 9, 10).

**Figure 1:**
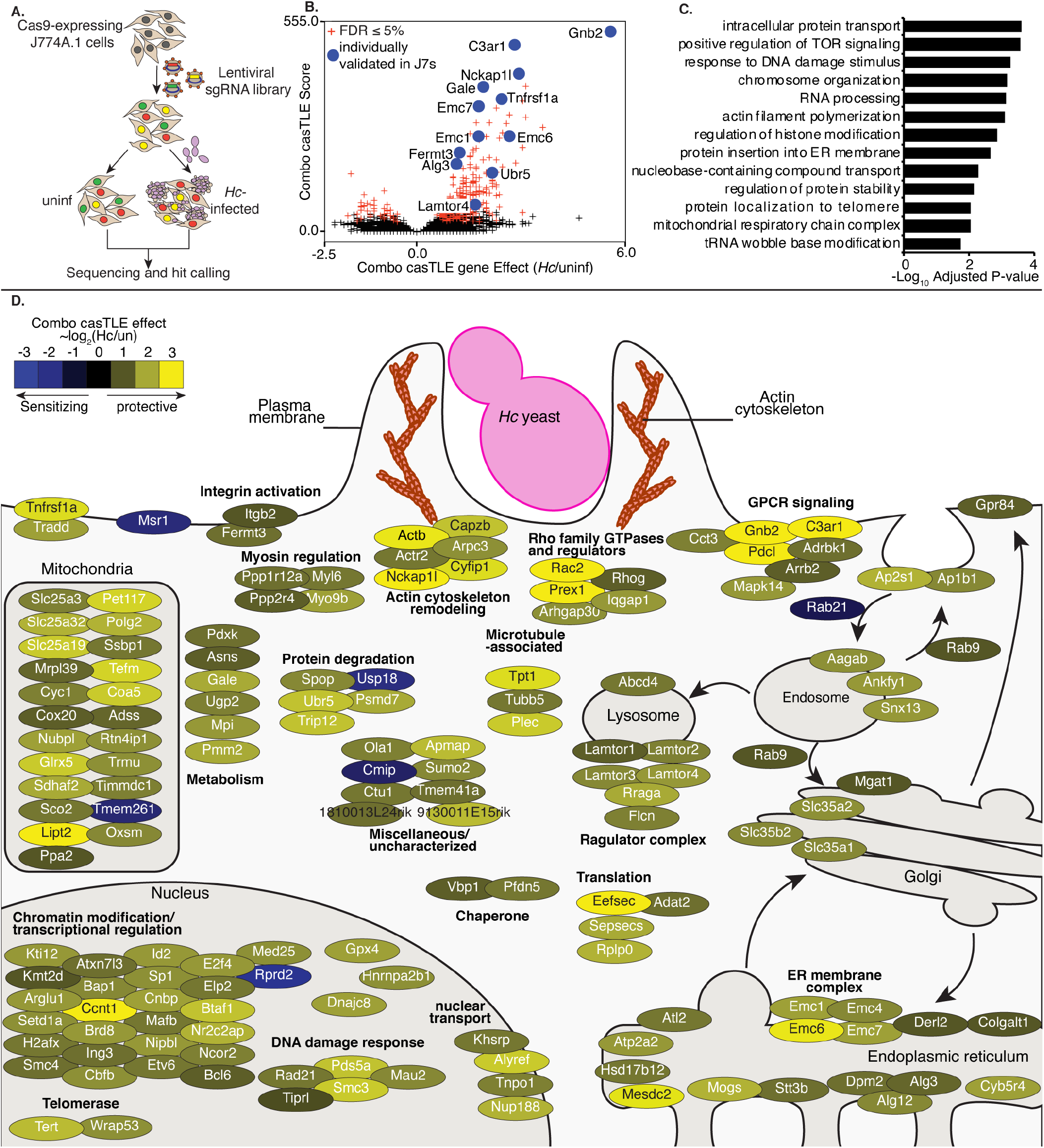
A pooled CRISPR screen identifies genes required for macrophage susceptibility to infection with *Hc*. **A.** Diagram of screen approach. Cas9-expressing J774A.1 macrophage-like cells were transduced with a library of sgRNAs, challenged with Ura5-deficient *Hc* yeast, and subjected to 2-3 pulses of uracil treatment followed by recovery. sgRNAs amplified from *Hc*-infected and uninfected cells were deep-sequenced, and sequences were analyzed to identify guides that became enriched or depleted in the *Hc*-infected pool relative to the uninfected pool. **B.** Volcano plot showing the confidence score (casTLE score) versus the effect size (casTLE effect) for all genes. Genes that pass the 5% FDR cutoff are colored red, and genes individually validated in J774A.1 cells are labelled and colored in blue. **C.** Adjusted P-values for selected GO biological process annotations enriched in the screen hits. **D.** The 150 highest-scoring genes identified in the screen grouped based on their annotated function and localization in a cell, functional categories or complexes of genes are noted. Genes are colored according to their gene effect estimate, where yellow indicates enrichment in the *Hc* infected pool and blue indicates depletion.

To create our knockout libraries, we first generated a clonal J774A.1 cell-line with high constitutive Cas9 activity (Fig. S1B). We then transduced these Cas9-expressing J774A.1 cells with pooled lentiviral sgRNAs. We used a previously designed CRISPR-Cas9 sgRNA library, which targets 23,000 protein-coding mouse genes with 10 sgRNAs/gene. The genome-wide library is split into 20 sub-libraries, each of which covers 500-1500 genes and includes 750 negative control sgRNAs (48). We screened each sub-library separately, covering a total of 16,782 genes. These cells were infected, in duplicate, with *Hc*, or were left uninfected and passaged throughout the course of the screen (Fig. 1A). To improve the sensitivity of our screen, we used a strain of *Hc* with a mutation in the *URA5* gene (*Hc ura5Δ*) which cannot grow in media without uracil supplementation (49). This strain does not lyse J774A.1 cells in the absence of exogenous uracil, and host cells that survive the initial round of lysis can be recovered by washing the monolayer and incubating in media without uracil supplementation (Fig. S1C-E), thereby allowing enrichment of resistant host cells. We infected the J774A.1 pools with *Hc ura5Δ* and performed 2-3 rounds of *Hc*-mediated lysis in the presence of uracil followed by uracil removal and recovery (see table S2 for sub-library specific details). We pulsed the uninfected cells with uracil during passaging to match the *Hc* infection. The sgRNAs in the final pools were deep-sequenced to determine the enrichment of guides following challenge with *Hc*. We employed the Cas9 high-throughput maximum-likelihood estimator (casTLE) algorithm (50) to estimate the effect of knocking out a gene on susceptibility to *Hc* (caSTLE effect) based on the enrichment of guides targeting each gene in the screen compared to the enrichment of negative control sgRNAs. We additionally analyzed uninfected cells at the beginning and the end of passaging using the casTLE algorithm, and we were able to verify that guides targeting genes previously annotated as essential (50) dropped out of the pool during passaging (Fig. S1H).

We identified 361 genes whose deletion modulated macrophage susceptibility to *Hc* infection at a 5% false-discovery rate (Fig. 1B). Confidence scores between screen replicates were moderately correlated (Fig. S1F). Disruption of 322 of these genes conferred protection against *Hc* (combo casTLE effect >0), and disruption of 39 conferred greater susceptibility to infection (combo casTLE effect <0) (Fig. 1B). We noticed that the protective hits include genes known to be required for macrophage phagocytosis, such as members of the SCAR/WAVE and ARP2/3 complexes (Fig. 1C-D). Such regulators have been well-studied for their role in phagocytosis and chemotaxis (14, 51, 52). Similarly, we identified *Itgb2* (CD18), which encodes the β-subunit of CR3 that has been previously shown to facilitate recognition and phagocytosis of *Hc* (16, 19), and *Fermt3*, which promotes activation of integrins (53) (Fig. 1D).

Of note, we identified a number of pathways and complexes among the resistance-promoting hits that have not been previously implicated in *Hc* interaction with macrophages (Fig. 1D), such as the ragulator complex, glycosylation enzymes, protein degradation machinery, mitochondrial respiration genes, solute transporters, and the ER membrane complex (EMC). The ragulator complex promotes nutrient stress sensing (54), and the EMC facilitates the folding of transmembrane proteins with multiple membrane-spanning regions (55, 56). The highest scoring protective hits include a group of genes (*Gnb2*, *Pdcl*, AP-1 subunits, AP-2 subunits, *Arrb2*) that regulate G-protein coupled receptor (GPCR) signaling and receptor trafficking following GPCR engagement (Fig. 1D) (57, 58). The hit identified with the second-highest confidence score was the gene encoding the GPCR C3a receptor 1 (*C3ar1/C3aR*) (Fig. 1D). Histograms demonstrating the enrichment of negative control sgRNAs and sgRNAs targeting *Gnb2* and *C3ar* in the *Hc-*infected pool are shown in Fig. S1G. We went on to investigate whether these factors play a role in macrophage phagocytosis of *Hc*.

### Identification of genes required for phagocytosis of yeast in J774A.1 macrophage-like cells and primary macrophages

We selected 16 high-confidence hits to individually validate in J774A.1 macrophage-like cells, including two genes, SCAR/WAVE subunit *Nckap1l*, and *Itgb2*, which were expected to play a role in macrophage phagocytosis of *Hc*. We prioritized genes that would shed light on novel aspects of macrophage interactions with *Hc* and that did not appear to strongly inhibit macrophage replication. We chose the three top-performing guides, based on enrichment or depletion in the screen, for further validation.

To verify susceptibility/resistance phenotypes in J774A.1 cells, we mixed GFP+, CRISPR-knockout (KO) cells with Cas9-expressing unlabeled cells, infected one pool of this mixture with *Hc*, and in parallel passaged the uninfected pool. Following one round of lysis and recovery, the pools were then harvested, and the proportion of GFP-expressing cells was measured via flow cytometry (Fig. 2A). The ratio of GFP+ cells in the *Hc*-infected compared to the uninfected pool demonstrated whether targeting a specific gene conferred a fitness advantage (>1) or disadvantage (<1) to macrophages during co-culture with *Hc*. Of the 16 genes tested, 13 conferred a fitness advantage during *Hc* infection when disrupted, including *Gnb2*, *C3ar*, ER membrane complex subunits *Emc1, Emc6,* and *Emc7*, and ubiquitin ligase *Ubr5* (Fig. 2B). As positive controls, we included knockouts of SCAR/WAVE subunit *Nckap1l*, and the β-2 integrin subunit of CR3, *Itgb2* (Fig. 2B). The only susceptibility-promoting hit that we tested, *Rab21*, did not promote increased susceptibility to *Hc* infection when disrupted (Fig. 2B).

**Figure 2:**
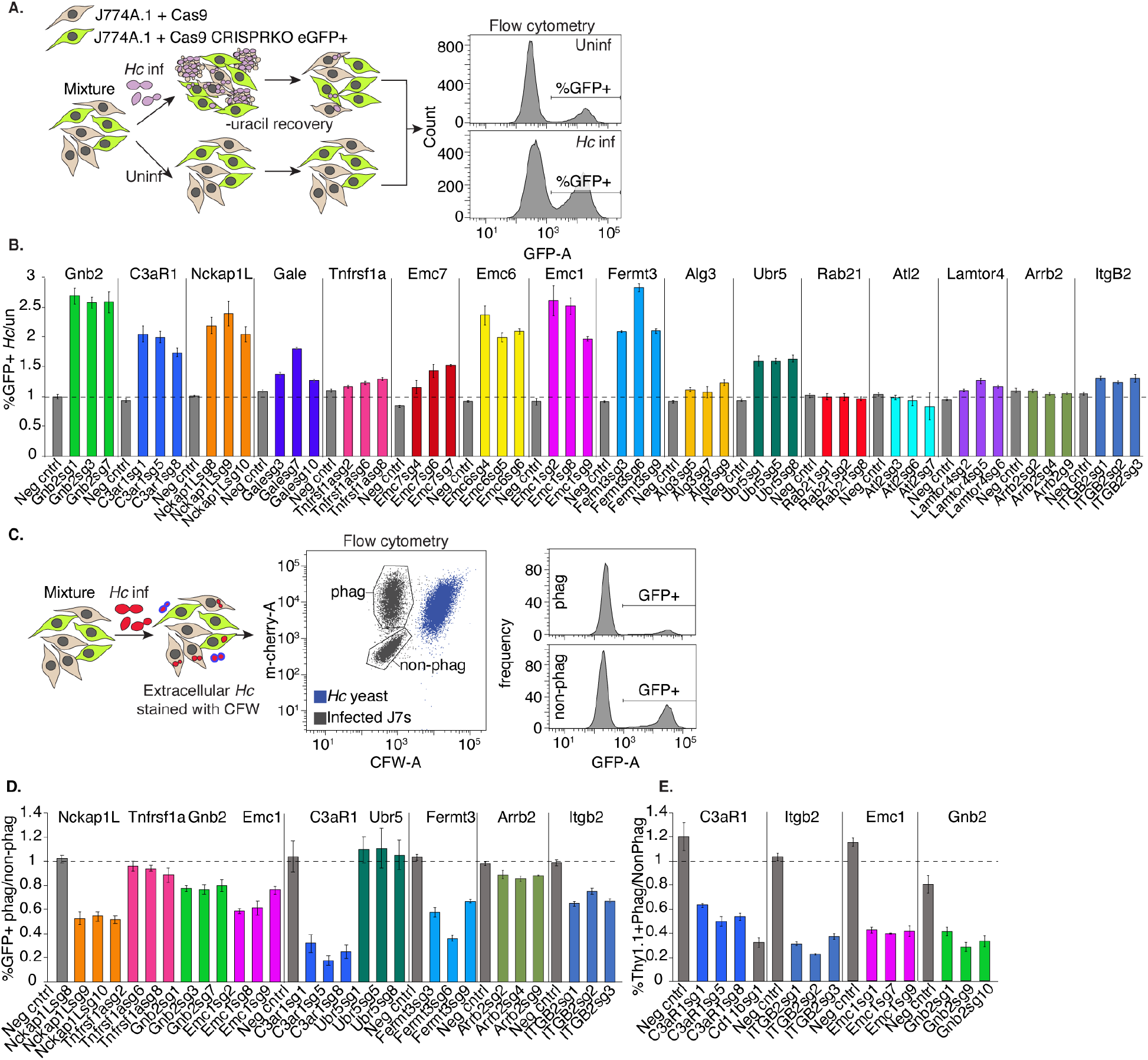
Identification of genes required for phagocytosis of yeast in J774A.1 cells and primary macrophages. **A.** Diagram of approach used to individually validate the role of a gene in macrophage susceptibility to *Hc* infection. A mixture of WT (GFP-) and CRISPRKO (GFP+) J774A.1 cells with or were challenged with *Hc* yeast in the presence of uracil, and allowed to recover. Uninfected cells from the same mixture were passaged in parallel, and the percentage of mutant cells in the *Hc* infected pools was compared to that of the uninfected pools via flow cytometry (n=3 biological replicates). **B.** Enrichment of gene-targeting guides in the *Hc* infected pool relative to the control pool, compared to that of non-targeting guides. **C.** Diagram of approach for determining the role of a gene in phagocytosis of *Hc.* A mixture WT (GFP-) and CRISPRKO (GFP+) J774A.1 cells were infected with mCherry-expressing *Hc* yeast. Non-internalized yeasts were excluded using calcofluor white staining. Flow cytometry was used to determine the representation of mutant cells in the phagocytic compared to the non-phagocytic populations (n=3). **D.** Identification of genes required for phagocytosis of yeast in J774A.1 cells using GFP expression to measure enrichment of sgRNA-expressing cells. **E.** Validation of gene involvement in BMDM phagocytosis of yeast using CRISPRKO BMDMs (Thy1.1+). A mixture of transduced (Thy1.1+) and untransduced (Thy1.1-) BMDMs were similarly infected with yeast and stained with calcofluor white and a Thy1.1 antibody to determine the representation of mutants in the phagocytic and non-phagocytic populations (n=3 biological replicates).

Next, we tested whether these genes play a role in macrophage phagocytosis of *Hc* yeast. To this end, we mixed GFP+, CRISPR-targeted cells with unlabeled, Cas9-expressing cells, infected the mixture with mCherry-expressing *Hc* yeast, and stained the cells with calcofluor white (CFW) to distinguish between intracellular and extracellular yeast. We used flow cytometry to measure the representation of GFP+ cells in the phagocytic compared to the non-phagocytic population (Fig. 2C). As expected, targeting of *Nckap1l, Fermt3,* and *Itgb2* led to decreased *Hc* phagocytosis in J774A.1 cells (Fig. 2D). Additionally, we found that knockout of *Emc1, Gnb2, C3ar1,* and *Arrb2* decreased phagocytosis of *Hc* (Fig. 2D).

Although J774A.1 cells recapitulate many important features of primary macrophages, including phagocytosis, they also differ in characteristics such as gene expression regulation (59). Therefore, we attempted to reproduce our findings from J774A.1 cells in bone marrow-derived macrophages (BMDMs) using CRISPR-Cas9-mediated gene disruption. We mixed CRISPR-knockout, Thy1.1+ BMDMs with WT, unlabeled BMDMs, infected the mixture with *Hc* yeast, and assessed phagocytosis as described above. We quantified the proportion of Thy1.1+ cells in the phagocytic compared to the non-phagocytic populations to determine whether the targeted genes promoted BMDM phagocytosis of *Hc* yeast. The four genes that we tested, GPCR *C3ar1*, integrin subunit *Itgb2*, ER membrane complex *Emc1*, and G*β* subunit *Gnb2*, were also required for efficient phagocytosis of *Hc* yeast by BMDMs (Fig. 2E).

### C3aR signaling plays a role in macrophage phagocytosis of fungi

Since a role for C3aR in phagocytosis of fungi had not previously been defined, we were intrigued by the result that this receptor is required for efficient phagocytosis of *Hc* by J774.1 cells and BMDMs. C3aR is a GPCR that recognizes the complement C3 cleavage product, anaphylatoxin C3a, and signals through G*α*i (29), which is sensitive to pertussis toxin-mediated ADP-ribosylation (60). We further investigated the role of C3aR and GPCR signaling in macrophage phagocytosis of fungi. We generated BMDMs from *C3ar*-/- mice (61) and age-matched WT mice. We then infected these macrophages with several species of pathogenic fungi including *Hc* yeast expressing mCherry (Fig. 3A-B, G-I)*, Candida albicans* (*Ca*) yeast detected with a fluorescent antibody (Fig. 3C), and *Coccidioides posadasii* arthroconidia labeled with FITC (*Cp*) (Fig. 3D), and determined the extent of phagocytosis over time. We also tested phagocytosis of FITC-labeled zymosan (Fig. 3B), a cell-wall extract of *Saccharomyces cerevisiae*. We used calcofluor white staining to distinguish between intracellular and extracellular fungi. We observed that C3aR was required for efficient phagocytosis of all three species of fungal pathogens, in addition to zymosan, suggesting an important general role for C3aR in macrophage capture and phagocytosis of fungi (Fig. 3A-D). The involvement of C3aR did not require fungal viability, as C3aR was equally important for phagocytosis of both live and killed *Hc* yeast, as well as zymosan (Fig. 3A-B). The phagocytosis defect was not due to a defect in CD11b or CD18 surface expression in *C3ar-/-* BMDMs (Fig. S2).

**Figure 3:**
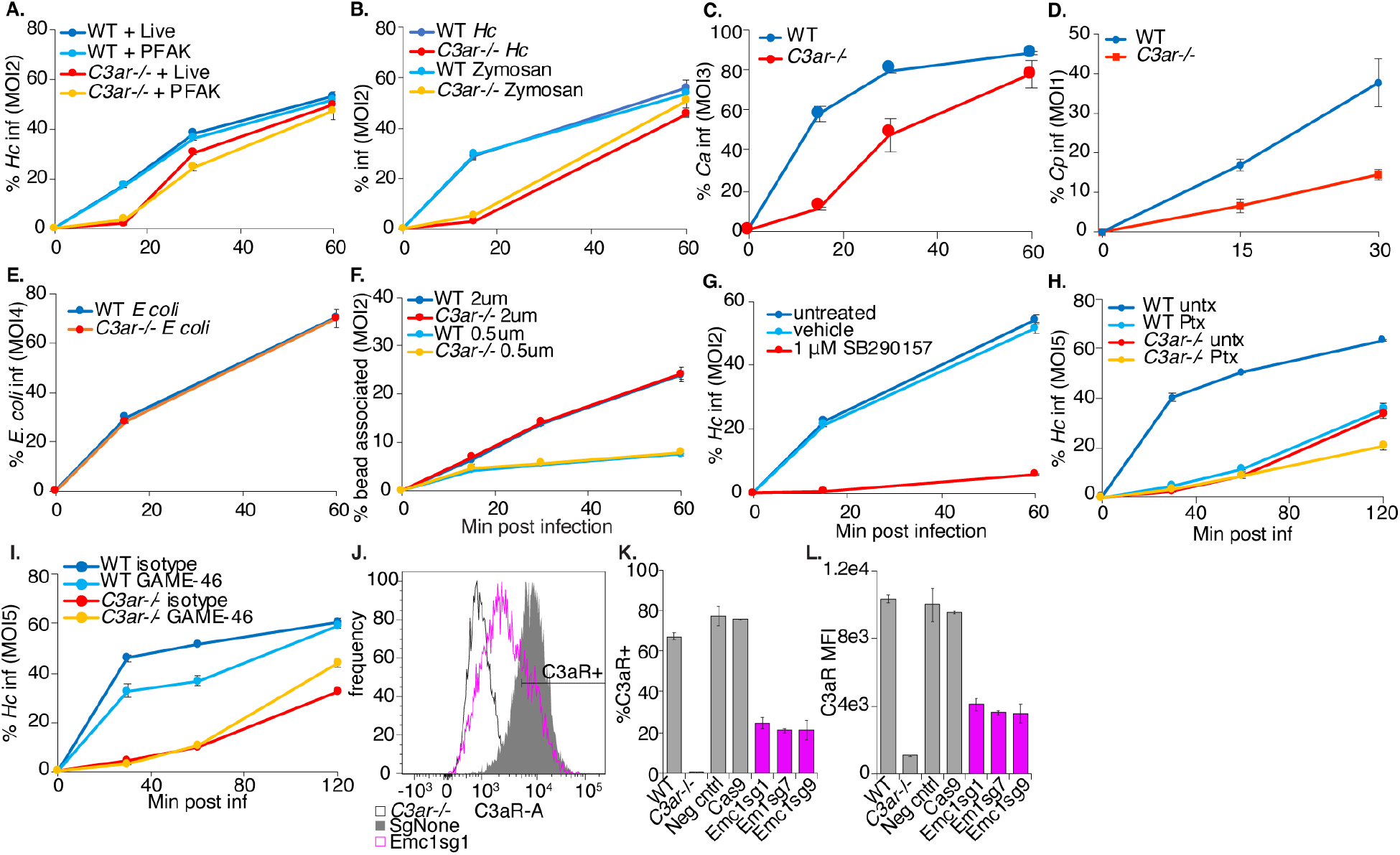
C3aR signaling plays a role in macrophage phagocytosis of fungi. **A.** WT and *C3ar-/-* BMDMs were infected with live and PFA-killed mCherry-expressing *Hc* yeast, and the phagocytosis rate was monitored over-time using flow-cytometry (n=3 biological replicates). **B.** WT and *C3ar-/-* BMDMs were infected with FITC-labelled zymosan or mCherry-expressing *Hc* and the phagocytosis rate infected cells was monitored using flow cytometry (n=3 biological replicates). **C.** BMDMs were infected with *Candida albicans* (*Ca*). Cells were imaged using confocal microscopy to quantify phagocytosis (n=2 biological replicates, >350 cells/replicate). CFW staining was used to exclude extracellular *Ca.* **D.** BMDMs were infected with FITC-labelled *Coccidioides posadasii* (*Cp*) arthroconidia, and extracellular conidia were labelled with calcofluor white. BMDM infection rates were determined using confocal microscopy (n=3 biological replicates, 200-400 cells/rep). **E.** BMDMs were infected with FITC-labelled *E. coli* bioparticles and the *E. coli-*association with BMDMs was monitored via flow cytometry (n=2 biological replicates). **F.** BMDMs were infected with 2 μm or 0.5 μm red fluorescent latex beads, and the rate of BMDM association with the beads was measured using flow cytometry (n=3 biological replicates). **G.** BMDMs were treated with a C3aR antagonist (1 μM SB290157) and infected with *Hc* yeast. Phagocytosis was measured using flow cytometry (n=3 biological replicates). **H.** BMDMs were pre-treated for 2 h with 1 μg/mL pertussis toxin (Ptx), which inhibits Gαi, and infected with *Hc* (n=3 biological replicates). **I.** BMDMs were pre-treated for 90 min with 10 μg/mL CD18 blocking antibody (GAME-46) and infected with *Hc* yeast (n=3 biological replicates) Phagocytosis was measured using flow cytometry. Emc1 is required for C3aR expression in BMDMs (**J-L**). **J.** *Emc1* CRISPRKO BMDMs and control sgRNA transduced BMDMs were stained with an anti-C3aR antibody, and C3aR levels were measured via flow cytometry (n=2 biological replicates). **K.** Histogram of C3aR levels in control and *Emc1* CRISPRKO BMDMs. **L.** Frequency of C3aR+ cells in the indicated BMDMs. **M.** The mean fluorescence intensity (MFI) of the C3aR signal in the indicated BMDMs.

C3aR has been previously implicated in macrophage uptake of certain, though not all, bacterial pathogens (62, 63), and in microglial phagocytosis of several substrates (64–66). To investigate whether the requirement of C3aR for phagocytosis extends to other types of particles that can be taken up by macrophages, we measured the capture of uncoated latex beads and FITC-labelled *E. coli* K12 in WT and *C3ar-/-* BMDMs (Fig. 3E-F). We found that C3aR was not required for uptake of *E. coli* (Fig. 3E) or latex beads (Fig. 3F), suggesting that C3aR does not play a general role in phagocytosis.

To further validate the contribution of C3aR to phagocytosis, we treated macrophages with a specific non-peptide antagonist of C3aR, SB290157 (67) five minutes before challenge with *Hc* (Fig. 3G). We found that the C3aR antagonist was able to inhibit macrophage phagocytosis of *Hc*, suggesting an acute role for C3aR in macrophage phagocytosis of fungi. C3aR signaling is dependent on pertussis toxin-sensitive G*α*i (29), inhibition of which interferes with macrophage phagocytosis of Zymosan particles (68). We assessed whether G*α*i inhibition by pre-treatment of macrophages with pertussis toxin (Ptx) would impact macrophage phagocytosis of *Hc* yeast, and whether Ptx treatment would synergize with C3aR deficiency. We found that Ptx pre-treatment inhibited macrophage phagocytosis of *Hc* (Fig. 3H). Ptx treatment strongly phenocopies the phagocytosis defect in *C3ar-/-* BMDMs, and Ptx treatment modestly inhibits phagocytosis in *C3ar-/-* BMDMs (Fig. 3H). These findings show that C3aR-dependent Gαi activation promotes phagocytosis, although Gαi activation by other receptors, and C3aR coupling to a different Gα subunit, may play a minor role in *Hc* phagocytosis (Fig. 3H). We also investigated whether C3aR interacts with CR3 to promote phagocytosis by treating BMDMs with a CD18 blocking antibody (GAME-46) previously used to block CR3(16) (Fig. 3I). WT BMDMs treated with the CD18 inhibitor had a modest defect in phagocytosis of *Hc*, and treatment of *C3ar-/-* BMDMs with the inhibitor did not further block phagocytosis of *Hc*, suggesting that CR3 participates in phagocytosis downstream of C3aR (Fig. 3I).

We further reasoned that Emc1 may indirectly promote phagocytosis due to its role in stabilization of proteins with multiple transmembrane helices (56), such as C3aR. To test this hypothesis, we measured C3aR expression in *Emc1* CRISPRKO BMDMs (Fig. 3J-L). We saw a dramatic decrease in C3aR expression in *Emc1*-targeted BMDMs compared to untransduced or control-targeted BMDMs (Fig. 3J-L), suggesting that the EMC facilitates the proper folding and biosynthesis of GPCRs, such as C3aR, in macrophages. In contrast, *Emc1* CRISPRKO BMDMs did not show reduced surface expression of CD18 or CD11b (Fig. S2), verifying that the EMC may not be as critical for proper folding of single-pass transmembrane proteins like integrins.

Since phagocytosis of *Hc* is delayed in *C3ar-/-* BMDMs, we expected lysis of infected BMDMs to show a corresponding delay. As expected, we found that *C3ar-/-* BMDMs were slightly less susceptible to *Hc*-mediated lysis, as measured by an established assay (9, 69) (Fig. S3). Analysis of *Hc* colony forming units (CFUs) indicated that *C3ar-/-* macrophages were infected with fewer *Hc* yeast at the start of the experiment, and *Hc* yeasts did not have a major intracellular growth defect in the mutant macrophages (Fig. S3). We did not observe a difference in *Hc*-induced TNFα secretion in the absence of C3aR, suggesting that C3aR does not affect macrophage cytokine release in response to *Hc* (Fig. S4).

### Serum C3 promotes complement opsonization and macrophage phagocytosis of *Hc* yeast

Since the canonical ligand for C3aR is C3a derived from C3 processing, we investigated whether serum represented a source of C3 that would react with *Hc* to generate C3a and promote phagocytosis. Macrophage infections discussed up to this point were conducted in the presence of 10% (for J774A.1 cells) or 20% (for BMDMs) heat-inactivated fetal bovine serum (FBS), which was not previously thought to be a robust source of complement.

We infected WT and *C3ar-/-* BMDMs with mCherry+ *Hc* and FITC-labelled zymosan in the presence or absence of 20% heat-inactivated FBS, and monitored phagocytosis by flow cytometry. Surprisingly, we found that FBS promoted macrophage phagocytosis of *Hc*, and to a lesser extent zymosan, in a C3aR-dependent manner (Fig. 4A). Even after 2h of co-culture, we did not observe efficient phagocytosis of *Hc* by BMDMs in serum-free media (Fig. 4B). Phagocytosis of zymosan by BMDMs in serum-free media was more efficient than that of *Hc*, and was not dependent on C3aR (Fig. 4A), as expected due to the role of Dectin 1-mediated recognition of β-glucans in non-opsonic macrophage recognition of zymosan (38). The low level of *Hc* phagocytosis in serum-free media was also C3aR-independent. The ability of FBS to stimulate phagocytosis is not lot-dependent, as FBS from different lots and manufacturers promoted macrophage phagocytosis of *Hc* in a C3aR-dependent manner (Fig. S5). To assess whether FBS was promoting phagocytosis by opsonization of the yeast, we tested whether pre-incubation with FBS would be sufficient to stimulate phagocytosis of *Hc* in serum-free media (Fig. 4B). We found that pre-incubation in FBS did not promote phagocytosis of *Hc* or zymosan (Fig. 4B), suggesting either that FBS does not facilitate phagocytosis by opsonization, or that opsonization is labile. We also determined that incubating *Hc* with BMDM conditioned media containing FBS did not promote macrophage phagocytosis of *Hc* (Fig. S6B), suggesting that BMDMs do not secrete a missing factor that would restore FBS-mediated opsonization of *Hc*.

**Figure 4:**
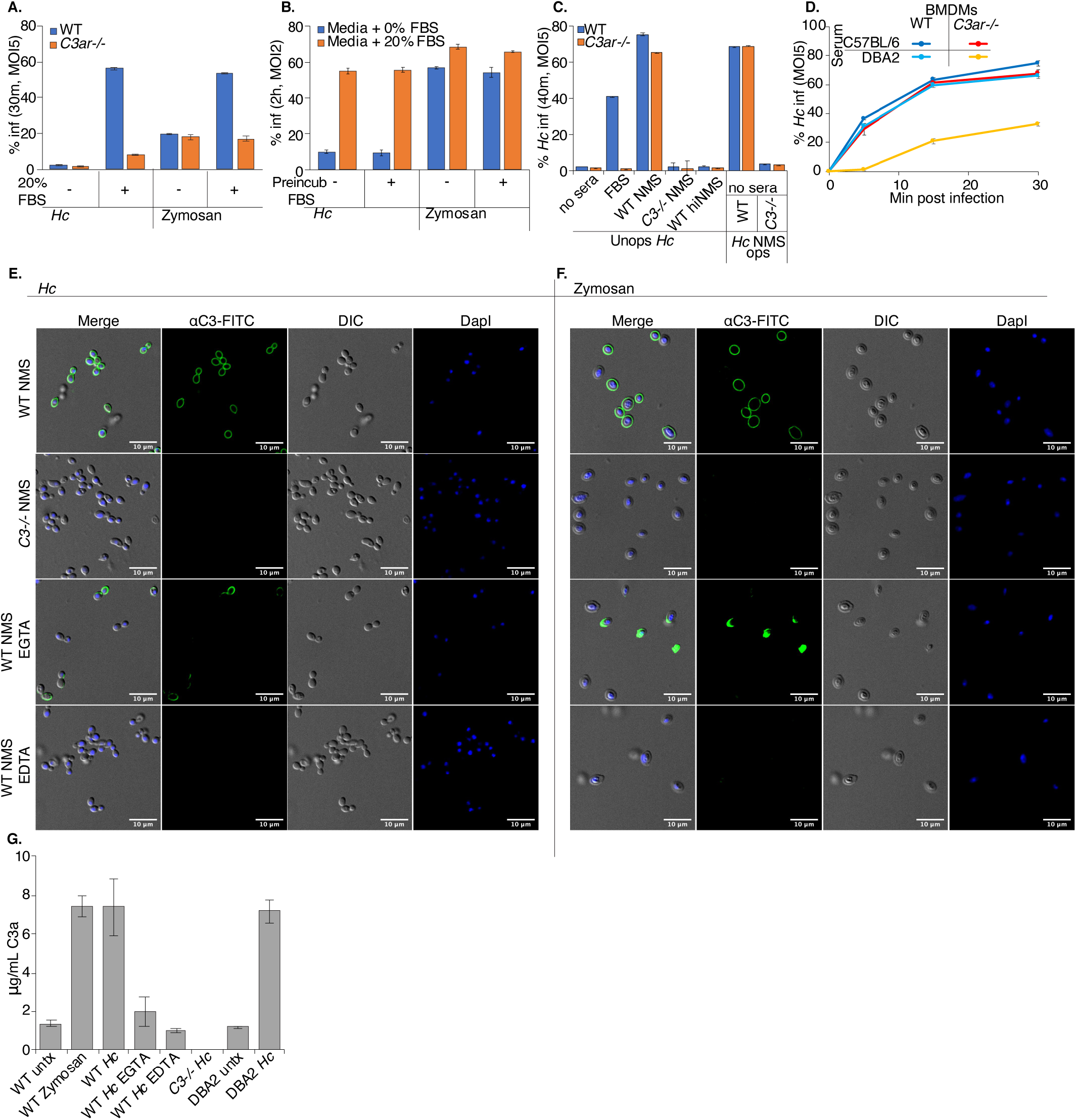
Serum C3 promotes complement opsonization and macrophage phagocytosis of *Hc* yeast. **A.** FBS stimulates macrophage phagocytosis of fungi in a C3aR-dependent manner. BMDMs were infected with mCherry-expressing *Hc* or FITC-labelled zymosan in the presence or absence of 20% heat-treated FBS (FBS). Phagocytosis was assessed via flow cytometry (n=3 biological replicates). **B.** FBS does not promote macrophage phagocytosis of *Hc* via opsonization. *Hc* and zymosan particles were pre-incubated with 10% heat-treated FBS for 30 min at 37°C, washed, and used to infect BMDMs. Phagocytosis was measured using flow cytometry (n=2 biological replicates). **C.** Normal mouse serum (NMS) stimulates BMDM phagocytosis of fungi in a C3-dependenent manner. BMDMs were infected with *Hc* yeast in serum-free media or media supplemented with 5% FBS, 5% NMS from WT mice, 5% NMS from C3-/- mice, or 5% heat-inactivated NMS (hiNMS) from WT mice. BMDMs in serum-free media were also infected with *Hc* opsonized with 10% WT or C3-/- NMS. Phagocytosis was measured as described above (n=3 biological replicates). **D.** C5-deficient serum promotes macrophage phagocytosis of *Hc* in a C3aR-dependant manner. BMDMs were infected with *Hc* yeast in media supplemented with 5% NMS from C57BL/6 mice or DBA2 (C5-deficient) mice. Phagocytosis was measured as described above (n=2 biological replicates). **E-F**. Mouse serum promotes complement opsonization of yeast via multiple pathways. *Hc* (**E**) or Zymosan (**F**) were incubated in PBS with 10% sera from WT or *C3-/-* mice. 10 mM EGTA or EDTA were added to the reactions to chelate Ca^2+^ or Mg^2+^, respectively. Yeast were stained with FITC conjugated anti-mouse C3, and imaged using confocal microscopy (representative slices are shown from 2 biological replicates). **G.** Incubation of *Hc* with mouse serum leads to C3a release. Supernatants were harvested following incubation of mouse serum with *Hc* or zymosan, and mouse C3a levels were measured by ELISA.

To establish a role for serum-derived C3 in macrophage recognition of *Hc*, we compared phagocytosis of *Hc* in media supplemented with no serum, FBS, or serum collected from WT or *C3-/-* C57BL/6 mice (WT NMS or *C3-/-* NMS) (Fig. 4C). We found that mouse serum promoted macrophage phagocytosis of *Hc* in a C3-dependent manner that was sensitive to heat inactivation (Fig. 4C). Surprisingly, the ability of mouse serum to stimulate phagocytosis of *Hc* was not dependent on C3aR (Fig. 4C), suggesting an additional C3aR-independent, C3-dependent mechanism of phagocytosis. Since C5 can be activated downstream of C3, leading to the release of the potent chemoattractant C5a (21), we reasoned that serum from C57BL/6 mice might stimulate phagocytosis via C5. C5a release and recognition by C5aR would then stimulate phagocytosis and compensate for C3aR-deficiency. To test this, we supplemented the media with serum from DBA2 mice, which have low levels of serum C5, but normal levels of C3 (70). We found that *C3ar-/-* BMDMs were defective at phagocytosis of *Hc* in media supplemented with DBA2 (C5-deficient), but not C57BL/6 (C5-sufficient) serum (Fig. 4D), suggesting that C5a in C57BL/6 serum acts redundantly with C3a to promote macrophage phagocytosis of *Hc*.

To confirm that incubating mouse serum with *Hc* yeast would promote opsonization with C3, as previously described (71), and C3a release, we incubated *Hc* yeast and zymosan (positive control) with mouse serum, visualized C3 deposition using immunofluorescence confocal microscopy (Fig. 4E-F) and measured C3a levels in the supernatant by ELISA (Fig. 4G). We observed robust C3 staining of *Hc* upon incubation with WT serum, and no C3 deposition after incubation with C3-/- sera (Fig. 4E). We also found that incubating WT C57BL/6 and DBA2 serum with *Hc* increased C3a levels in the supernatant (Fig. 4G), suggesting C3a release. To inhibit the classical/lectin pathways, or all activation pathways, we added EGTA or EDTA, respectively, to the indicated reactions. We did not observe C3 deposition or C3a release when Mg++ was chelated with EDTA (Fig. 4E). We also saw C3 deposition, although with lower efficiency and with a less uniform distribution around the yeast cell-wall, and lower levels of C3a release, in the presence of EGTA (Fig. 4E-G). These results confirm that *Hc* can activate the alternative complement pathway, as previously suggested (34). Due to the increased efficiency and uniformity of C3 deposition on yeast and the increased C3a release found in the absence of EGTA, we suggest that the classical or lectin pathways also contribute to C3 opsonization of *Hc* yeast. We did not find evidence of C3 deposition on the cell-surface following incubation of *Hc* with FBS or BMDM conditioned media containing FBS (Fig. S6A). To demonstrate that complement opsonization by mouse serum promotes macrophage phagocytosis of *Hc* yeast, we infected BMDMs in serum-free media with *Hc* opsonized by WT or *C3-/-* mouse serum. We found that opsonization with WT mouse serum, but not *C3-/-* serum, is sufficient to promote phagocytosis of *Hc* in serum-free media in a C3aR-independent manner, suggesting direct recognition of opsonized yeasts by CR3 (Fig. 4C). This activity was blocked by EDTA and moderately inhibited by EGTA, suggesting contribution of both the classical/lectin and alternative pathways to phagocytosis stimulation through opsonization (Fig. S6B).

Active complement C3 can also be secreted by macrophages (30–32). We measured the release of C3 into culture supernatants by ELISA, and found that *Hc* infection did stimulate a modest macrophage secretion of C3 (Fig. S7A). However, we did not observe a phagocytosis defect when we infected *C3-/-* BMDMs with *Hc* or zymosan in the presence of FBS (Fig. S7B), suggesting that macrophage-derived C3 is not playing a major role in macrophage phagocytosis of *Hc* in our assay.

### C3aR localizes to the early *Hc-*containing phagosome

We next analyzed C3aR localization during macrophage phagocytosis of *Hc* and latex beads, whose uptake does not depend on C3aR. These experiments were conducted in media supplemented with 20% FBS. Localization to the *Hc* containing phagosome would implicate C3aR directly in fungal capture or phagocytic cup formation. Immunofluorescence confocal microscopy confirmed that C3aR is localized at the plasma membrane (Fig. S8). We observed C3aR localization to the *Hc*-containing phagosomes at 5- and 10-minutes post-infection, and with a lower frequency at 30 minutes post-infection (Fig. 5A). Examples of C3aR-positive phagosomes are indicated by white arrows in the images. In contrast, we did not observe C3aR-positive bead-containing phagosomes at the same frequency (Fig. 5B), suggesting that C3aR localizes specifically to the *Hc*-containing phagosome, and not to latex bead-containing phagosomes.

**Figure 5:**
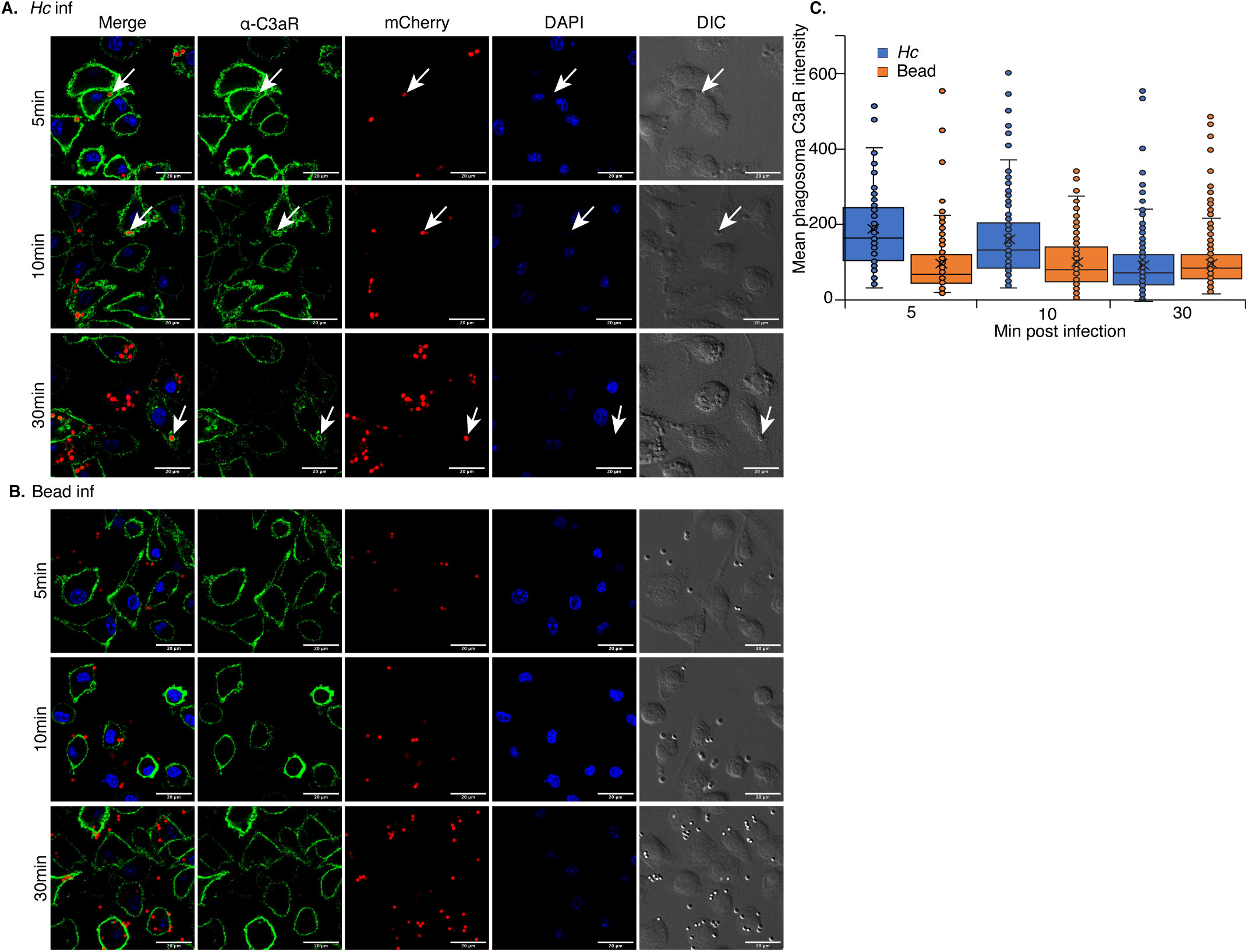
C3aR localizes to the early *Hc*-containing phagosome. C3aR localizes to *Hc*-containing phagosomes (**A**) to a greater extent than latex bead-containing phagosomes (**B**). BMDMs were infected with the indicated particles (MOI=5, n=2 biological replicates per time point). Cells were then stained with a C3aR-specific antibody and imaged using optical sectioning with a confocal microscope. Representative images from a single slice are shown. **C.** The mean fluorescence intensity of C3aR in the particle-containing phagosomes was quantified using ImageJ (N>91 phagosomes).

To quantify C3aR localization to the *Hc* or bead-containing phagosome, we used imageJ to measure the mean intensity over background of the C3aR signal surrounding the *Hc* or bead particle. Our analysis revealed that *Hc*-containing phagosomes display significantly higher C3aR enrichment than bead-containing phagosomes (T-test, P-value <0.05) at 5- and 10-minutes post-infection, but not at 30 minutes post-infection as the phagosomes mature (Fig. 5C).

### C3aR promotes the formation of actin-rich protrusions that facilitate capture of *Hc* yeast

Since C3a is a chemoattractant for macrophages, we investigated the role of macrophage migration in the C3aR-dependent capture of *Hc* yeast. Although macrophages did undergo chemotaxis towards *Hc* in trans-well migration assays, migration was not dependent on FBS or C3aR (Fig. S9). We also were not able to rescue the phagocytosis of *Hc* by *C3ar-/-* macrophages when the likelihood of *Hc*-macrophage interaction was increased by centrifugation of *Hc* onto the monolayer, or an extended pre-incubation on ice (Fig. S10). These experiments suggest that C3aR involvement in macrophage phagocytosis of *Hc* is not due to its role in facilitating long-range migration of macrophages towards yeast. However, these studies do not rule out a role for C3aR-dependent control of short-range chemotaxis in macrophage capture of *Hc* yeast.

To investigate this possibility, we generated J774A.1 cells that express Lifeact-mEGFP, a probe that specifically labels F-actin (72), and performed live imaging of J774A.1 macrophages during co-culture with mCherry-labelled yeast in the presence of a C3aR antagonist (SB290157) or a vehicle control using confocal microscopy (Fig. 6). These movies show macrophages extending actin-rich membrane protrusions in the direction of nearby *Hc* that promote rapid *Hc* capture and engulfment (example time series shown in Fig. 6A). In contrast, C3aR antagonist-treated macrophages show much slower capture of *Hc* yeast, and fail to rapidly form such actin-rich directed membrane protrusions (Fig. 6B). Membrane protrusions of macrophages that eventually capture *Hc* yeast were tracked and analyzed (73) (Fig. 6C-E). Treatment with the C3aR antagonist dramatically slowed capture of *Hc* yeast, as demonstrated by the lower phagocytosis rate and the lower mean velocity of the tracked protrusions (Fig. 6D). Finally, the movement of the antagonist-treated membrane protrusions was less directional, as demonstrated by the lower outreach ratio of the membrane protrusions (Fig. 6E).

**Figure 6:**
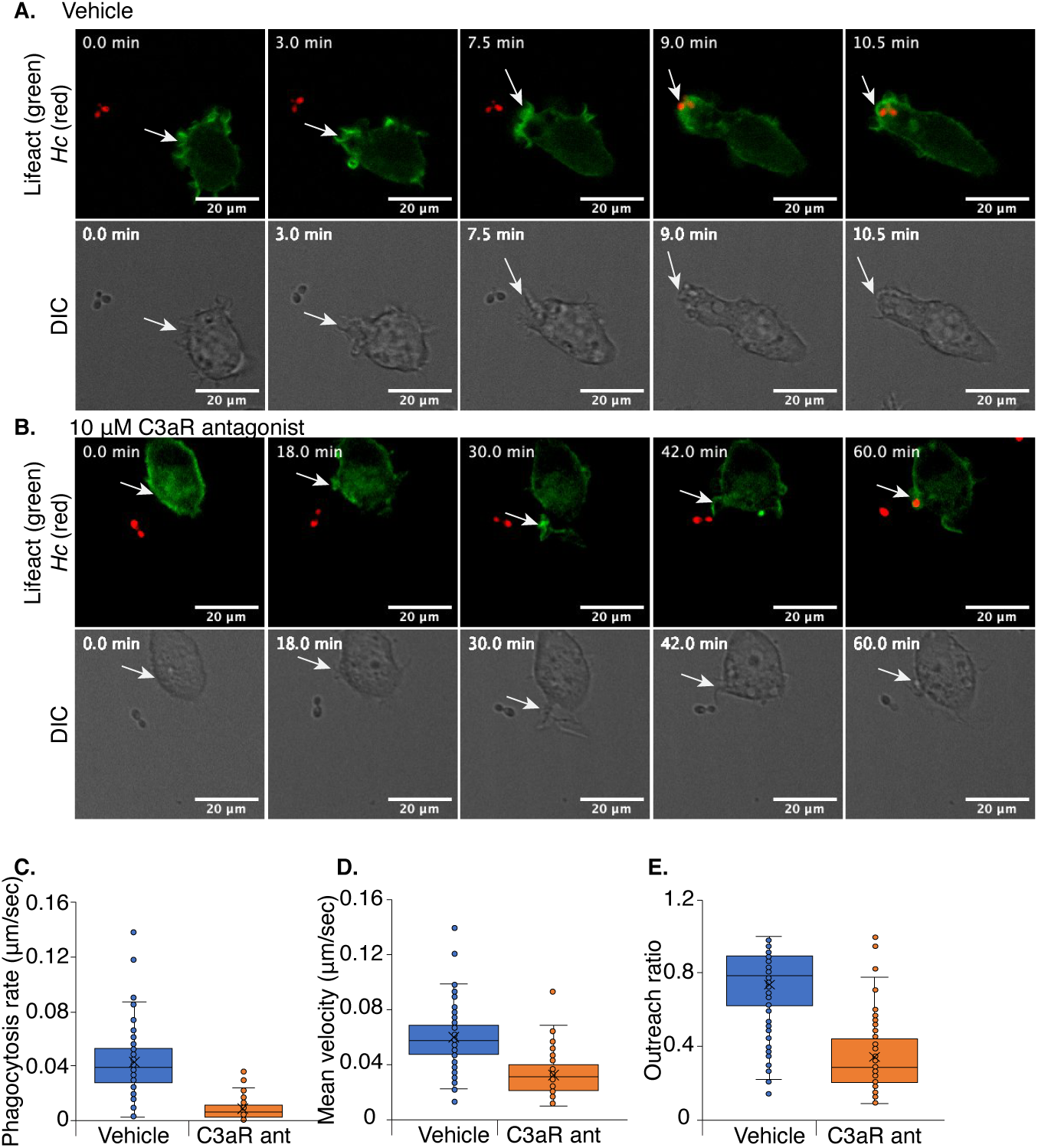
C3aR promotes the formation of actin-rich protrusions that facilitate capture of *Hc* yeast. J774A.1 cells were engineered to express Lifeact-mEGFP to label F-actin, co-cultured with mCherry-expressing *Hc* yeast, and subjected to live-cell confocal microscopy in a temperature- and-CO_2_ controlled chamber in media supplemented with 10% FBS. Cells were treated with a C3aR antagonist (10 μM SB290157) or a vehicle control. **A.** Representative images from a confocal time series showing a macrophage extending an F-actin-rich protrusion towards an mCherry expressing *Hc* yeast, followed by phagocytosis and formation of an actin-rich phagosome. The corresponding DIC images are shown below. **B.** A similar time series of macrophages treated with SB290157 showing a failure to initiate formation of a membrane protrusion and much slower capture of *Hc* yeast. The movement of membrane structures that successfully caputured yeast were analyzed using MtrackJ to quantify the behaviors of these structures (**C-E**), including the phagocytosis rate, quantified as the time required for the macrophage to successfully engulf the yeast divided by the distance of the yeast to the macrophage at the start of the series (**C**), the mean velocity of the membrane structure closest to the yeast (**D**), and the outreach ratio quantified as the max displacement of the track divided by the length of the track (**E**). These metrics demonstrate that macrophages treated with the C3aR antagonist are defective at the extension of membrane protrusions in the direction of *Hc* yeast that facilitate phagocytosis.

Live imaging experiments showed that C3aR facilitates the directional movement of actin-rich membrane protrusions towards *Hc* yeast that facilitate rapid phagocytosis. This behavior likely requires a C3a gradient that diffuses away from the *Hc* yeast following complement cleavage at the fungal surface. Consistent with this idea, the addition of recombinant C3a to BMDMs in the absence of a gradient was not sufficient to stimulate macrophage phagocytosis of *Hc* in serum-free media (Fig. S11).

## Discussion

We report a large-scale CRISPR-Cas9 screen conducted in macrophage-like cells challenged with *Hc* yeast. 361 genes emerged as high-confidence modifiers of macrophage susceptibility to *Hc*-mediated killing, vastly expanding our knowledge of the gene networks that underpin macrophage interaction with this important pathogen. Validation of top hits revealed an under-appreciated role for GPCR signaling through C3aR in macrophage phagocytosis of fungi. These results are particularly intriguing for *Histoplasma*, which is an intracellular fungal pathogen that thrives within the macrophage phagosome. Therefore, elucidating the molecular events that govern *Histoplasma* phagocytosis is particularly important for understanding *Hc* pathogenesis. Given that C3aR strongly enhances the efficiency of *Hc* phagocytosis, it may be a key host factor that promotes the intracellular lifestyle of this organism.

It was previously established that macrophage phagocytosis of *Hc* is not dependent on β-glucan recognition by Dectin-1 (16), and that *Hc* utilizes a number of mechanisms to minimize exposure of β-glucan on the cell surface (17, 18). In contrast, CR3 has been previously implicated in non-opsonic uptake of *Hc* (16, 19). Our work uncovers the important role of C3aR as a pattern recognition receptor for *Hc* and other fungi, potentially collaborating with CR3 to facilitate uptake of pathogenic yeasts that shield β-glucan from recognition by Dectin-1 (described in Fig. 7). We also discovered that C3aR-dependent phagocytosis requires serum, and that only mouse serum that was replete with C3 could stimulate phagocytosis, suggesting that a gradient of C3a emanating from the fungal surface might be critical for the phagocytic activity of C3aR as discussed below. Since *Hc* cannot engage Dectin-1, there is little phagocytosis of *Hc* in the absence of serum. In contrast, the residual serum-independent, C3aR-independent phagocytosis of zymosan may be due to Dectin-1. Given that *Hc* is introduced to the host via inhalation, and since complement activity is present in the bronchoalveolar fluid (22, 23), innate immune recognition of *Hc* likely occurs in the context of complement activation *in vivo*.

**Figure 7:**
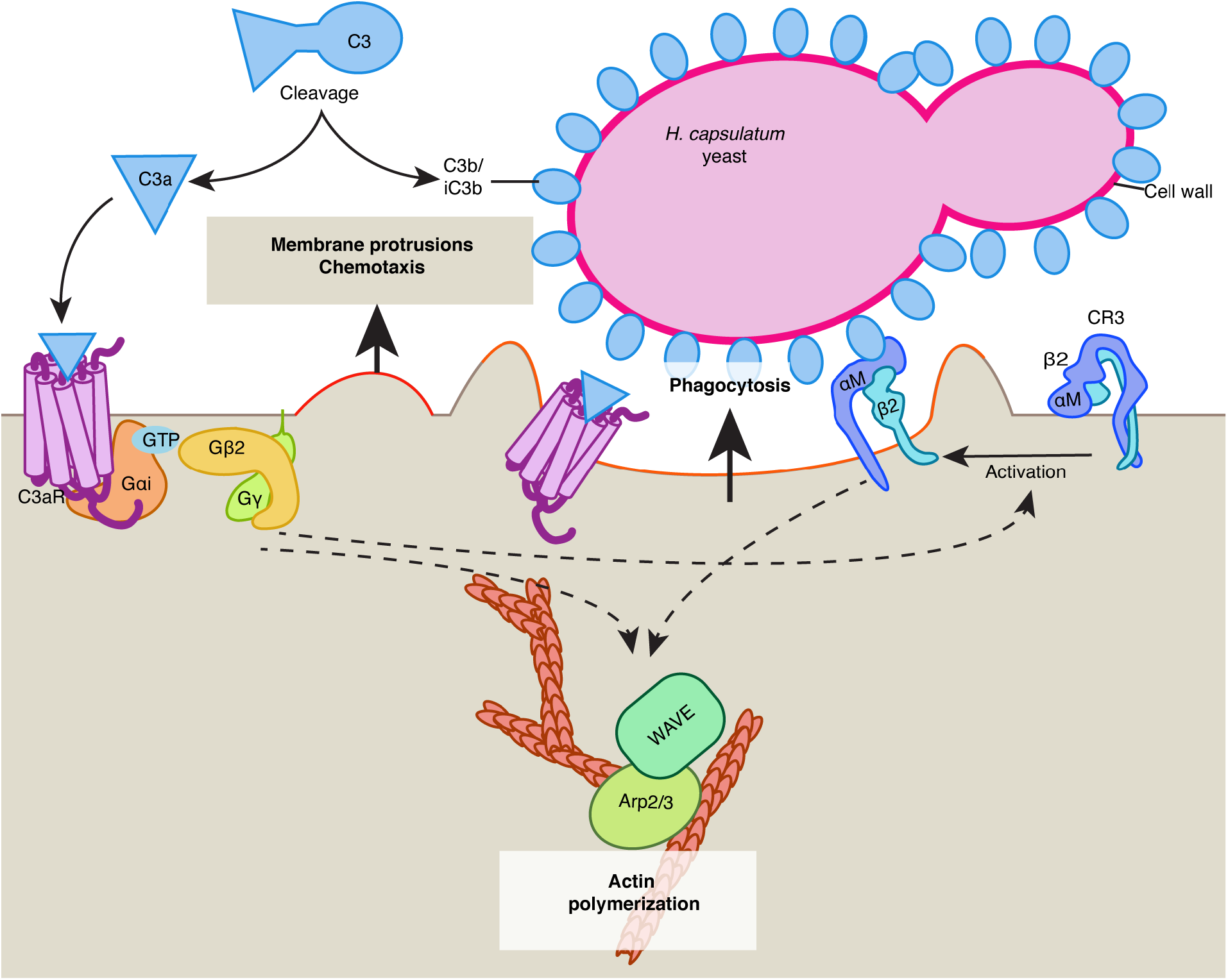
Model for the role of complement and C3aR in macrophage recognition of *Hc* yeast. We propose the following model for the role of complement and C3aR in macrophage recognition of *Hc*: C3, derived from serum, reacts with the *Hc* cell-wall, leading to C3b/iC3b deposition on the cell-wall, and release of C3a, which diffuses away from the yeast surface leading to a concentration gradient emanating from the yeast cell-wall. C3a activates C3aR, which signals through Gαi and Gβ2 to promote the formation and directional movement of actin-rich membrane protrusions, and possibly to promote activation or increased motility of the integrin receptor CR3. Active CR3 can then recognize C3b/iC3b or other features of the *Hc* cell-wall. C3aR and/or CR3 activation then coordinates actin polymerization and phagocytic cup formation by regulating the activity of actin polymerization regulators Arp2/3 and SCAR/WAVE.

The vast majority of genes identified in the screen were resistance-promoting hits, which may reflect limitations in the pooled screening approach, or the efficiency at which *Hc* evades macrophage defenses (in other words, it is challenging to increase macrophage sensitivity to *Hc*). Within these hits, we identified genes with previously described involvement in phagocytosis and *Hc* recognition, which validates our approach and is consistent with the requirement for *Hc* uptake to trigger the process of macrophage lysis. Our screen also revealed a role for GPCR signaling in *Hc*-host interactions. In addition to *C3ar*, the highest scoring protective hits included a set of genes that regulate signaling and receptor trafficking following GPCR engagement (57, 58). We validated that several of these genes, including *C3ar1, Gnb2,* and *Arrb2,* facilitate macrophage phagocytosis of *Hc.* While G-protein coupled receptor (GPCR) signaling is traditionally thought to play a role in chemotaxis rather than phagocytosis (74, 75), several studies have implicated G-protein activity directly in cytoskeleton coordination during phagocytosis (68, 76, 77). Both chemotaxis and phagocytosis depend on precise regulation of the actin cytoskeleton, and signaling often converges on the same signaling cytoskeleton remodeling machinery (74). Additionally, previous studies have shown that the mobility and activity of phagocytosis receptors is increased at the leading edge of a cell (78), and that active probing of the local environment by macrophages is critical for efficient binding of targets (79), suggesting strong coordination between chemotaxis and phagocytosis. We also identify the ER membrane complex, which facilitates the folding of transmembrane proteins with multiple membrane-spanning regions (55, 56). We show that *Emc1* promotes macrophage phagocytosis of *Hc*, and is required for surface expression of C3aR, but not CR3 subunits. Thus, we propose that the EMC indirectly participates in phagocytosis due to its role in folding receptors such as C3aR.

Other genes and complexes identified in this screen may play important roles in *Hc* interaction with macrophages. To uncover the nature of their involvement will require further study. These include the ragulator complex, which activates mTORC1 upon nutrient deprivation and regulates autophagic flux that can be critical for defense against intracellular pathogens (54). This complex also has been found in screens for phagocytosis regulators (51), and has been shown to modulate phagocytic flux in microglia (80). Other hits may affect *Hc*-macrophage interactions through indirect means, or promote nutrient acquisition and intracellular replication within the phagosome. We also identified ubiquitin ligases such as *Ubr5* and *Trip12*, which regulate histone ubiquitylation upon DNA damage (81). *Ubr5* has also been shown to down-regulate TLR signaling (82). Validation in macrophage-like cells demonstrates that *Ubr5* is required for *Hc-*induced lysis, but not macrophage phagocytosis of *Hc*, suggesting that Ubr5 promotes intracellular replication of *Hc* or macrophage cell-death.

Since the identification of C3aR as a phagocytic receptor was intriguing, we further characterized its role in macrophage phagocytosis of *Hc* and other targets. While we found that C3aR was required for phagocytosis of several species of fungi, C3aR did not play a general role in phagocytosis, as *C3ar-/-* macrophages were not defective in uptake of *E. coli* or latex beads. Previous studies have demonstrated that C3aR promotes phagocytosis of damaged neurons (83), myelin particles (66) and protein aggregates (64). C3aR has also been implicated in macrophage phagocytosis of uropathogenic *E. coli* (63), but not *Pseudomonas aeruginosa* (62). Further study is needed to determine the shared characteristics of particles that require C3aR for optimal phagocytosis, such as particle size, reactivity with complement, or other biochemical properties. Nonetheless, the identification of C3aR as an important phagocytic receptor for fungi implies that it may play a critical role in host defense to fungal pathogens. More study is needed to determine whether C3aR affects host susceptibility to fungal pathogens or modulates the immune response to fungi. Such study will be essential to determining the therapeutic benefit of targeting complement or C3aR in the treatment of invasive fungal infections.

We found that heat-inactivated fetal bovine serum (FBS) added to the macrophage media promoted fungal phagocytosis in C3ar-dependent but opsonization-independent manner. This suggests that FBS promotes phagocytosis predominantly by generating C3a that activates C3aR, although the mechanism by which FBS-derived C3a might play a role independent of C3b opsonization is unclear. We also showed that mouse serum was able to stimulate macrophage phagocytosis of *Hc* in a C3-dependent manner, and that C3 opsonization of *Hc* promoted macrophage phagocytosis, consistent with studies showing a role for C3b/C3bi in recognition of fungal pathogens (35). Surprisingly, the ability of serum from C57BL/6 mice to promote macrophage phagocytosis of *Hc* was not dependent on C3aR. We demonstrated that C5a-C5aR signaling compensated for C3aR deficiency, since macrophage phagocytosis of *Hc* in the presence of serum from DBA2 mice, which are C5-deficient, was dependent on C3aR. This is not surprising given that C5a and C5aR have been previously implicated in innate immune recognition of fungi (42, 43, 84).

To investigate the role of C3aR in macrophage phagocytosis of *Hc,* we demonstrated that C3aR localizes to the *Hc* containing phagosome at early time-points during infection. Localization of C3aR to the phagosome suggests direct involvement of C3aR in *Hc* recognition or cytoskeleton remodeling. Alternatively, C3aR might not directly participate in phagosome formation, but display enrichment at the plasma membrane sites that participate in *Hc* phagocytosis.

Finally, live imaging of actin dynamics in macrophages during *Hc* infection revealed that C3aR promotes the directional movement of actin-rich membrane protrusions that aid in the capture of *Hc* yeast. This is consistent with the ability of C3a to promote chemotaxis of innate immune cells including macrophages (27), and the role of G-protein signaling in activating cytoskeleton remodeling at the leading edge and the phagocytic cup (68, 76, 77). We did not find strong evidence that C3aR promotes chemotaxis towards *Hc* yeast in trans-well assays, and we were not able to restore phagocytosis in *C3ar-/-* macrophages by forcing contact between macrophages and *Hc,* suggesting that C3aR participates in short-distance rather than long-distance migration during fungal phagocytosis. C3aR may also promote optimal phagocytosis by spatially coordinating receptor mobility (78) or activation (58) at the leading edge. We believe that a gradient of C3a diffusing away from the *Hc* surface is critical this activity, as the uniform distribution of recombinant C3a alone was not sufficient to stimulate macrophage phagocytosis in the absence of serum. More investigation is needed to untangle the precise mechanism by which the C3a-C3aR pathway contributes to *Hc* recognition.

## Materials and Methods

### Strains and culture conditions

J774A.1 cells (ATCC) were cultured in Dulbecco’s modified Eagle’s medium high glucose (DMEM, UCSF media production) with 10% heat-inactivated fetal bovine serum (FBS; Corning or Atlanta), penicillin and streptomycin (pen/strep, UCSF media production). Cells were passaged by detaching with a disposable cell scraper. HEK293T cells (ATCC) were cultured in DMEM with 10% FBS and pen/strep. Platinum-E (Plat-E) retroviral packaging cells (CellBioLabs) were a gift from Jason Cyster (UCSF) and were maintained in DMEM supplemented with 10% FBS, pen/strep, glutamine, and 10mM HEPES (UCSF media production). Plat-E and HEK293T cells were passaged by detaching cells using 0.05% Trypsin-EDTA (UCSF media production). WT C57BL/6J (stock 000664), Rosa26-Cas9 (stock 26179), *C3ar-/-* (stock 33904), *C3-/-* (stock 29661), and DBA2/J (stock 000671) mice were obtained from Jackson Laboratories and bred in the UCSF mouse barrier facility. Bone marrow from 6-to 8-week-old female mice was isolated from femurs and tibias, and differentiated into bone marrow-derived macrophages (BMDMs) by culturing in BMM (bone marrow macrophage media) + 10mM HEPES as described previously(85). BMM contains 10% CMG-conditioned media and 20% FBS. Mammalian cells were frozen in complete media supplemented with 10% DMSO and 50% FBS, and stored in liquid nitrogen. *Histoplasma capsulatum* (*Hc*) strain G217B (ATCC 26032) and G217B *ura5Δ* were kind gifts from William Goldman (University of North Carolina, Chapel Hill). mCherry-expressing *Hc* was generated as described previously(86). The *Hc cpb1* mutant strain, G217Bura5Δcbp1::T-DNA with a Ura5-containing episomal vector, was generated previously(7, 9). *Hc* cultures were grown on *Histoplasma* macrophage medium (HMM) agarose plates or in liquid HMM on an orbital shaker as previously described(87). Mammalian cells and *Hc* cultures were maintained in humidified tissue-culture incubators at 37°C with 5% CO_2_. *Hc* was grown on HMM-agar plates (supplemented with 0.175 mg/mL uracil to grow *Hc ura5Δ*) for 1-2 weeks, and passaged in 1:25 HMM liquid culture every-other day for five days to obtain logarithmic-phase *Hc* yeast-cultures (OD_600_=5-7). Yeast cells were collected, resuspended in Ca^++^ and Mg^++^-free D-PBS (D-PBS), sonicated for 3 seconds on setting 2 using a Fisher Scientific Sonic Dismembrator Model 100, and counted using a hemocytometer. *Hc* yeast were adjusted to the appropriate concentration in D-PBS. For macrophage infections, *Hc* was added to the macrophage cultures, and allowed to settle onto the cells unless otherwise specified. *Candida albicans* (*Ca*) strain Sc5314 (ATCC MYA-2876) was a kind gift from Alexander Johnson (UCSF). *Ca* was grown on YEPD (2% peptone, 1% yeast extract, 2% glucose) agar or liquid media at 30°C. *Coccidioides posadasii* Silveira strain was a generous gift from Dr. Bridget Barker (Northern Arizona University). Coccidioides arthroconidia were obtained as previously described(88), by growing Coccidioides on 2xGYE (2% glucose 1% yeast extract) solid agar in flasks at 30°C for 4-6 weeks. At the time of collection, arthroconidia were dislodged with a cell scraper in PBS, filtered through miracloth to remove hyphal fragments, resuspended in PBS and stored at 4C for up to 6 months. Arthroconidia concentration was measured by counting arthroconidia on hemocytometer.

### Generation of stable J774A.1 cell-lines for CRISPRKO and live-cell imaging experiments

Gene-targeting sequences were cloned into the pMCB306 lentiguide-puro vector as previously described(89). Table S3 lists the targeting sequences used. The lentiviral Lifeact-monomeric eGFP-Blast vector was a kind gift from Diane Barber (UCSF). The Ef1a-Cas9-Blast lentiviral vector (pMCB393) was generated previously(48). To generate lentivirus particles, HEK293T cells were transfected using polyethylenimine (PEI) with second-generation (sgRNA, Lifeact) or third-generation (Ef1a-Cas9-Blast) packaging plasmids and the desired transfer plasmid. Lentivirus was 48- and 72-h later, and filtered through a 0.45 μm polyvinylidene fluoride (PVDF) or polyethersulfone (PES) filter (Millipore). Viruses were concentrated using the Lenti-X concentrator (Takara) according to the manufacturer’s instructions. Concentrated lentivirus (Cas9: 20X, lentiguide-puro: 1-2X, Lifeact: 5X) was added to J774A.1 cells for 12-24 h (with 8 μg/mL polybrene for Cas9), after which virus-containing media was removed and cells were grown in complete DMEM. Starting at 3 days post-transduction, cells were grown under selection with Blasticidin (2 μg/mL) or puromycin (2.5 μg/mL) for 3 days, and expanded without selection for at least 3 days or until the desired number of cells was obtained. To obtain clonal Cas9-expressing J774A.1 cells, live cells were harvested and single-cell sorted using a FACSAriaII cell-sorter into 96-well plates containing complete media supplemented with 50% sterile-filtered J774A.1 conditioned media, and expanded for 3 weeks. The Cas9 activity of the J7-Cas9 clones was determined following transduction with the lentiguide-puro-eGFP vector containing a GFP-targeting sgRNA, and measuring eGFP silencing after puromycin selection by flow cytometry. The J7-Cas9 clone with the highest eGFP-silencing activity was used to generate the pooled CRISPR libraries and individual CRISPRKO cell-lines. The efficiency of Cas9-mediated gene-targeting was assessed by PCR-amplifying the targeted locus in control and CRISPRKO cells, performing Sanger sequencing, and analyzing sequencing chromatograms using the TIDE webtool(90).

### Pooled CRISPR-Cas9 screens

We used pooled mouse sgRNA sub-libraries that were generated previously(48), some of which are available on Addgene (#1000000121-1000000130). Each library covers 500-1500 genes with 10 sgRNAs/gene and includes 750 negative control sgRNAs (375 non-targeting and 375 safe-targeting sgRNAs). We performed screens on all of the sub-libraries, except for the Mouse Unique sub-libraries, which contain mouse genes that do not have known orthologues in humans. Taken together, our screens covered 16,781 mouse genes. Lentivirus was generated by transfecting HEK293T cells seeded in 15 cm dishes with sgRNA plasmids and second-generation packaging plasmid as described previously(91). Lentivirus was harvested at 48- and 72-h post-transfection, filtered through 0.45 μm PES filters, pooled, then concentrated using the Lenti-X concentrator (Takara) according to the manufacturer’s instructions. J774A.1 cells stably expressing LentiCas9-Blast (generation described above) were incubated with 2X concentrated lentivirus for 24h at 1000X coverage in T-225 or T175 flasks for an MOI of 0.2-0.5 as determined by flow cytometry of mCherry expression at 3 days post-transduction. We then performed selection for transductants using puromycin (2.5 μg/mL) for 3 days until >90% of the cells were mCherry-positive by flow cytometry. Cells were allowed to recover from puromycin selection for three days before screening. Cells were split into two conditions, and screening was performed in duplicate. One condition was infected with *Hc ura5Δ* and subjected to 2-3 pulses of uracil to initiate fungal growth and macrophage lysis (see Table S2 for details specific to each sub-library). J774A.1 CRISPRKO libraries, seeded at 1000X library coverage in T-225 or T-150 flasks, were infected with *Hc ura5Δ* at a multiplicity of infection (MOI) of 5 yeast/macrophage. Yeast were allowed to settle onto the monolayer and incubated for 2 h. The cells were washed once with D-PBS to remove extracellular yeast, and incubated in the presence of 0.35 mg/mL uracil for 2 d until ∼50% of the monolayer was cleared. Then, the monolayer was washed 3X with D-PBS to remove dead macrophages and extracellular yeast, and incubated for 2-5 days in complete media without uracil to allow the monolayer to recover. Then, uracil was re-introduced to the culture media for 1-2 d to re-initiate fungal growth and lysis. The addition and removal of uracil was performed 1-2 times depending on the speed at which the monolayer recovered. Uninfected cells were passaged in parallel every 2 d by detaching adherent cells with a cell-scraper, counting using a hemocytometer, and re-seeding into new flasks at 1000X coverage. Uninfected cells were pulsed with uracil during passaging to match the *Hc* infection. At the end of the screening period, cells were washed and harvested by detaching with a cell-scraper. Genomic DNA was extracted from the cells using the DNA blood midi or maxi kit according to the manufacturer’s instructions, with the inclusion of a brief centrifugation step after cell lysis to remove un-lysed *Hc* yeast before addition of ethanol and application to the column. Guide frequencies were quantified by PCR amplification and deep sequencing using an illumina NextSeq 500 as previously described(89).

### Analysis of CRISPR-Cas9 Screens

Sub-library screens were analyzed separately using casTLE version 1.0 as previously described(50). Briefly, the distribution of guides was compared between the uninfected and *Hc*-infected samples, and guide enrichments were calculated as log ratios between the infected and uninfected samples. A maximum likelihood estimator was used to estimate the effect size for each gene and the log-likelihood ratio (confidence score, or casTLE score) by comparing the distribution of the 10 gene-targeting guides to the distribution of negative control guides. An effect size of 1 roughly corresponds to one log2 fold change of the gene compared to the negative controls. P values were determined by permuting the gene-targeting guides in the screen and comparing to the distribution of negative controls using casTLE, and false discovery rate (FDR) thresholds for defining hits were calculated using the Benjamini-Hochberg procedure. We used a threshold of 5% FDR to define hits. Results from the separate sub-library screens were concatenated and visualized together using JavaTreeview(92). GO-biological process analysis was performed using Gorilla(93) using an un-ranked list of genes that passed the 5% FDR cutoff as the target list and all of the genes detected in the screen as the background list.

### Competitive fitness assays in J774A.1 cells

J774A.1-Cas9 (WT) cells were mixed with CRISPRKO J774A.1-Cas9 cells harboring the lentiguide-puro vector, which drives expression of a gene-targeting sgRNA and an eGFP marker (75% WT cells, 25% CRISPRKO cells). 3.5X10^5^ cells/well were seeded in tissue culture (TC)-treated 6-well plates. 12-24 h later, the cells were infected with *Hc ura5*Δ at an MOI=5, which was incubated with the monolayer for 2 h followed by a D-PBS wash step. The cells were incubated in complete media containing 0.35 μg/mL uracil for 2 d, until lysis of >50% of the monolayer was observed. Then cells were recovered by washing 3X with D-PBS, and incubating in complete media in the absence of uracil for 2 d. Uninfected cells were detached by scraping and passaged to prevent overcrowding, and were subjected to the same washing and media conditions as the *Hc*-infected cells. Following the recovery period, surviving cells were harvested and stained, and GFP-expression was analyzed by flow cytometry.

### Generation of CRISPR-knockout BMDMs

The pSIN MuLV sgRNA retroviral transfer plasmid (U6 guide tracer EF1a Thy1.1 P2A Neo) was a kind gift from Jason Cyster (UCSF). The sgRNA cloning site, U6 promoter, and selection marker of pSIN was replaced to match that of pMCB306 using the Gibson Assembly Cloning Kit (NEB) to generate the transfer plasmid (BAS2186) used for these studies. Gene-targeting sgRNA sequences (Table S3) were cloned into the vector as previously described for pMCB306(89). To generate viral particles for expression of sgRNAs, Plat-E retroviral packaging cells were transfected with the transfer plasmid in antibiotic-free complete DMEM. Virus was harvested at 48 h and 72 h post-transfection and filtered through a 0.45μm PES filter. Bone marrow from female 6-8-week-old Rosa26-Cas9 mice was isolated and cultured for 2 d in BMM as described above. Non-adherent bone marrow cells were harvested, and 2X10^6^ cells per well were infected with 2 mL fresh MuLV supernatant by centrifugation (2400 RPM, 2 h, RT) in 6-well non-TC-treated plates with 10 μg/mL polybrene. Viral supernatant was removed, and cells were incubated overnight in BMM. Both adherent and non-adherent bone marrow cells were infected with viral supernatant again as described above with the 72h viral harvest. 24h after the second viral spinfection, BMDMs were grown under puromycin selection (4 μg/mL) for 3 days, grown for an additional 3-5 days in BMM without puromycin, and harvested as previously described. Retroviral infection and selection were verified by Thy1.1 staining and flow cytometry. The efficiency of Cas9-mediated gene-targeting was assessed by PCR-amplifying the targeted locus in control and CRISPRKO cells, performing sanger sequencing, and analyzing sequencing chromatograms using the TIDE webtool(90).

### Competitive *Hc* phagocytosis assays

WT and CRISPRKO J774A.1 cells were mixed as described above (75% WT and 25% CRISPRKO), and seeded at 3X10^5^ cells/well in tissue-culture-treated 12-well plates and incubated for 12-24 h prior to infection. *Hc* yeast expressing mCherry were added to the monolayers at an MOI=2, and incubated for 1h at 37°C. Cells were then washed with ice-cold HBSS and harvested by pipetting the cells off of the well with HBSS. Similarly, Cas9-expressing BMDMs (WT) were mixed with Cas9-BMDMs transduced with a retroviral vector driving expression of a gene-targeting sgRNA (CRISPRKO) (75% WT and 25% CRISPRKO). BMDMs were added at 5X10^5^ cells/well to non-TC-treated 12-well plates in BMM for 12-24 h, then infected with mCherry-expressing *Hc* for 1 h in BMM. Phagocytosis and GFP or Thy1.1 expression was measured using flow cytometry.

### FITC labelling of Zymosan and *Coccidioides posadasii* arthroconidia

FITC-labelling was performed as described previously for *Hc* yeast(16). Briefly, Zymosan A (Sigma) was sonicated for 3 seconds on setting 2 using a Fisher Scientific Sonic Dismembrator Model 100, washed with 0.05 M sodium carbonate-bicarbonate buffer, and adjusted to 2X10^8^ particles/mL. *C. posadasii* arthroconidia were adjusted to 5X10^8^ conidia/mL, and washed in a sodium carbonate-bicarbonate buffer. Fungi were incubated with in 0.05M sodium carbonate-bicarbonate buffer (pH 9.5) with 0.16mg/mL FITC (Fisher, dissolved in DMSO at 5mg/mL) for 15 min at room temperature, protected from light, then washed twice with HBSA (HBSS + 0.25% BSA). Labelled zymosan was resuspended in D-PBS, counted using a hemocytometer, and frozen in single-use aliquots at −20°C. FITC-labelled arthroconidia were resuspended in PBS and counted on a hemocytometer. FITC-labelled arthroconidia were kept at 4°C and protected from light until used in phagocytosis experiments.

### Mouse serum collection

Mice were euthanized using CO_2_, and blood was collected by cardiac puncture. Blood was placed in a 1.5 mL Eppendorf tube and allowed to coagulate by incubation at room temperature for 1 h, and for an additional 30 min on ice. Coagulated blood was centrifuged, and the supernatant was harvested. Serum was used fresh or stored at −80°C in single-use aliquots.

### BMDM phagocytosis assays

BMDMs were thawed and seeded in BMM in12-well non-TC-treated plates at 5X10^5^ cells/well (flow cytometry) or onto ethanol-sterilized glass coverslips in TC-treated 24-well plates at 2X10^5^ cells/well (microscopy), and allowed to adhere for 12-24 h. Prior to infection, the cells were washed with D-PBS and fresh media was added. For some experiments, BMDMs were pre-treated with the following inhibitors or vehicle controls: pertussis toxin (Sigma, 1 μg/mL 2 h), SB290157 (Sigma, 1 μM 5 min), anti-CD18 clone GAME-46 or isotype control (BD, 10 μg/mL 90 min). To ensure a consistent MOI across BMDM harvests, prior to infection a well of BMDMs was harvested using dissociation buffer, and counted using a hemocytometer. The count was used to calculate the number of BMDMs that had adhered to the dish to determine the number of fungal cells or particles to add to the well for the appropriate MOI. For *Hc* phagocytosis assays, mCherry-expressing *Hc* yeast was added to the macrophage monolayer. For Zymosan phagocytosis assays, FITC-labelled zymosan was sonicated and added to the monolayer. Fluorescein-conjugated *E. coli* bioparticles (Invitrogen) were prepared according to the manufacturer’s instructions and added to macrophage monolayers at an MOI=4. Carboxylate-modified microspheres (2.0 μm and 0.5 μm) were sonicated and added to the macrophage monolayers at an MOI of 2. Phagocytosis of the above substrates was analyzed by flow cytometry. Logarithmic cultures of *Candida albicans* yeast grown in YEPD liquid media were harvested, washed 3X with D-PBS, counted, and added to macrophage monolayers on coverslips at an MOI of 3. Coverslips were washed 2X with DPBS and stained with 35 μg/mL calcofluor white for 1 min. Then, the coverslips were washed, fixed with 4% PFA at 37°C for 20 min. PFA was quenched by washing 3X with 100 mM glycine. Cells were permeabilized with 0.1% Triton-X-100 (5 min), and blocked with 1% BSA. Both intracellular and extracellular *C. albicans* yeast were detected by staining with a FITC-conjugated anti-*C. albicans* antibody (abcam, 1:1000) overnight 4°C. FITC-labelled *C. posadasii* arthroconidia were added to the coverslip an MOI of 1, then spun for 15 min at 550g to ensure contact between macrophages and arthroconidia. At the indicated times, the coverslips were washed and stained with 35 μg/mL CFW (2 min), washed once, fixed with 4% PFA, then washed with PBS. For *C. albicans* and *C. posadasii* experiments, Coverslips were mounted and imaged at 40X magnification. 16 fields along a grid were automatically selected. To determine the phagocytosis rate, the cell-counter plugin in FIJI was used to manually count the total number of macrophages and the number of macrophages with at least one intracellular fungus, determined by exclusion of CFW staining. The phagocytosis rate was calculated as the number of macrophages with at least one intracellular fungal particle divided by the total number of macrophages.

### Serum opsonization and analysis of C3 deposition by immunofluorescence microscopy

*Hc* or zymosan was incubated at 1X10^8^ particles/mL with 10% serum and the indicated chelators (10 mM EGTA or EDTA) in PBS for 30 min at 37°C. The yeast/particles were washed 2X with PBS, and co-cultured with BMDMs, or stained with a FITC-conjugated anti-mouse C3 antibody (MP Biomedicals, 1:200) for 1h at RT. Following staining, yeast/zymosan were washed 2X with PBS, and fixed with 4% PFA after spinning onto poly-L-Lysine-coated coverslips. Coverslips were washed 2X and imaged at 60X magnification to visualize mouse C3 deposition on the cell-wall.

### Analysis of C3aR localization by immunofluorescence microscopy

2X10^5^ BMDMs were seeded onto ETOH-sterilized glass coverslips in 24-well plates. Phagocytosis was synchronized by pre-incubating macrophage monolayers on ice, and centrifuging mCherry *Hc* yeast or fluorescent latex beads (MOI=5) at 4°C onto the monolayers, followed by incubation at 37°C for up to 30 min. Coverslips were washed with D-PBS and fixed with 4% PFA for 20min at RT. Coverslips were blocked with PBS + 5% FBS for 1 h at RT, and stained with an anti-mouse C3aR antibody (Clone 14D4, Hycult, 1:1000) overnight at 4°C in 5% FBS. Coverslips were washed with 5% FBS and stained with AlexaFluor-488-conjugated goat anti-rat IgG (Invitrogen, 1:500) for 1 h at RT. Coverslips were imaged at 60X magnification. Optical sectioning was performed to obtain Z-stacks (0.4 μm step-size, 5 μm thickness), and 6 fields were imaged per coverslip. To quantify C3aR localization to the *Hc* or bead-containing phagosome, we used imageJ to define the phagosome perimeter using thresholding and binary operations on the *Hc* or the bead channels. Then, we use the 3D ROI manager(94) plugin on imageJ to quantify the mean intensity of the C3aR signal within the phagosomal volume. We subtracted the background signal, measured on phagosomes in *C3ar-/-* BMDMs subjected to the same staining and analysis pipeline.

### Live cell-imaging and cell tracking

5X10^3^ Lifeact-meGFP-expressing J774A.1 macrophages (generated as described in “generation of stable cell-lines”) were seeded into 96-well glass-bottom plates (Cellvis) and allowed to adhere for 12-24 h. Culture media was replaced with fresh phenol-red-free DMEM containing 10% FBS with either a vehicle control (DMSO) or 10 μM C3aR antagonist (SB290157) and incubated for 5 min. mCherry-expressing were added to the cells at an MOI of 5, and centrifuged briefly (15 sec) to facilitate contact with the macrophages. Cells were imaged every 90 sec for 45 min at 20X magnification. An Okolab stagetop incubator with temperature and humidity control was used to maintain optimal conditions (37°C and 5% CO_2_). Four fields were imaged per duplicate well. Actin-rich (eGFP+) membrane protrusions of macrophages that capture *Hc* yeast were tracked manually using the imageJ plugin MtrackJ(95). Tracking was started at the membrane point closest to the *Hc* yeast when the yeast first appeared close to the location at which it was eventually captured. The position of the yeast was used as a reference. The track was terminated when the *Hc* was successfully engulfed (as visualized by formation of an actin collar around the *Hc* yeast), or when the imaging period terminated. The phagocytosis rate is reported as the distance from the J774A.1 cell-membrane to the *Hc* target at the start of tracking divided by the time elapsed until the yeast was successfully engulfed. The mean velocity (mean displacement/time across tracked points) and outreach ratio of the tracks (the max displacement/net displacement) were calculated as described(73).

### Confocal microscopy

For fixed imaging, coverslips were mounted onto slides using vectashield antifade mounting media, with or without DAPI (Vector labs) and sealed using nail polish. Fluorescence confocal microscopy was performed using a using a Nikon Ti-Eclipse inverted microscope with a Yokogawa spinning disk CSU-X1 and an Andor Clara CCD camera. Image analysis was performed using FIJI (ImageJ).

### Flow cytometry

BMDMs were washed and harvested using HBSS-based cell dissociation buffer (Thermo scientific) by incubating at 37°C for 10 min and pipetting with ice-cold HBSS. J774A.1 cells were washed with ice-cold HBSS, and harvested by spraying cells off of the well with ice-cold HBSS using a P1000 pipette. Cells were kept on ice and protected from light for subsequent steps. Cells were stained with fixable viability dye efluor450 (ebioscience; 1:1000) for competitive fitness assays and CD11b/CD18 staining, or fixable viability dye efluor780 (ebioscience; 1:500) for phagocytosis assays for 20 min. Cells were washed with FACS buffer (2% FBS and 5 mM EDTA in PBS) prior to staining with antibodies and/or Calcofluor White M2R (Sigma, 1 μg/mL) in FACS buffer for 15-20min. The following antibodies and dilutions were used: PerCP-conjugated anti-Thy1.1 (clone OX-7, biolegend, 1:100), PE, FITC, or AlexaFluor647-conjugated anti-CD11b antibody (clone M1/70, UCSF mAB core, 1:500 for BMDMs, 1:1000 for J774A.1), and AlexaFluor-647-conjuaged anti-CD18 antibody (M18/2, Biolegend, 1:100). Cells were washed with FACS buffer. For phagocytosis and competitive fitness assays, cells were washed with D-PBS, and fixed using BD stabilizing fixative for 15min, washed with D-PBS, and kept on ice prior to data acquisition. For indirect flow cytometry measurement of C3aR expression, 5X10^5^ BMDMs were fixed using BD stabilizing fixative (20min on ice). Cells were blocked for 20min with PBS5 (PBS+5% FBS) and stained with a C3aR antibody (Clone 14D4, Hycult, 1:500) in PBS5 for 20min on ice. Cells were washed with PBS5 and stained with APC-conjugated goat anti-rat IgG (Biolegend, 1:200) for 20 min. Cells were washed with PBS5, and resuspended in PBS for flow cytometry analysis. Flow cytometry acquisition was performed using a BD LSRII analyzer in the UCSF Parnassus Flow Core (RRID:SCR_018206). Analysis was performed using FlowJo v. 7 or 10. Where necessary compensation was performed with single-color controls using FlowJo.

### Trans-well migration assay

Cells and *Hc* were resuspended in migration media (DMEM with 0.5% fatty acid-free BSA, pen/strep, and 10 mM HEPES) or complete DMEM with 10% FBS. Inhibitors were added to both the well and the insert when used. 6.5 mm transwell permeable supports with 5 μm pore polycarbonate membranes (Costar) were used. Migration assays were performed in duplicate. 600 μL *Hc* G217B yeast was added at the indicated concentration to the well of a 24-well transwell plate. 2X10^5^ J774A.1 cells in 100 μL media were seeded into the transwell insert, and plates were incubated at 37°C with 5% CO_2_ for 3 h with minimal disturbance. Media was removed from the insert, which was dipped once in D-PBS, then placed in crystal violet stain (0.5% crystal violet, 20% methanol) for 10 min at RT. Supports were rinsed with dH_2_O, and a Q-tip was used to gently wipe cells off of the top of the membrane, and dried at RT. Membranes were mounted on slides, and 3 fields per membrane were imaged using a Leica DM 1000 microscope at 10X magnification. Cells in each microscopic field were counted manually using the cell counter plugin in FIJI.

### Cytotoxicity assays

7.5X10^4^ BMDMs were seeded per well of a 48-well plate and infected with *Hc* G217B at an MOI of 0.5 in the presence of pheno-red-free BMM. 1.875X10^4^ J774A.1 cells were seeded per well of a 48-well plate and infected with *Hc* at an MOI of 5 in phenol-red-free complete DMEM. Where indicated, media was supplemented with 0.35mg/mL uracil. To recover J774A.1 cells from infection with *Hc ura5Δ*, cells were washed with D-PBS, and grown in complete media that did not contain uracil for 3 days. Recovered cells were re-seeded and incubated with complete media with or without uracil supplementation. At the indicated time points, the amount of lactate dehydrogenase (LDH) in the culture supernatant was measured as described previously(96). Macrophage lysis is calculated as the percentage of total LDH from uninfected macrophages lysed in 1% Triton-X at the time of infection. Due to continued replication of macrophages during the experiment, the total LDH at later time points is greater than the total LDH from the first time point, resulting in an apparent lysis that is greater than 100%. To quantify cell depletion and recovery during infections of J774A.1 cells, we measured macrophage DNA remaining in the wells as previously described(10). Briefly, we washed the cells with D-PBS, added ddH_2_O to the well to lyse the macrophages, and measured the amount of macrophage DNA in the wells using the picoGreen DsDNA reagent (Invitrogen). Fluorescence intensities were measured using the quantitative plate read option on an Mx3000P QPCR machine (Agilent).

### Intracellular fungal growth assay

7.5X10^4^ BMDMs were seeded per well of a 48-well plate and infected in triplicate with *Hc* at an MOI of 0.5. At the indicated time points, culture supernatants were removed and 500 μL ddH_2_O was added. Macrophages were osmotically and mechanically lysed, and plated on HMM agarose at the appropriate dilutions as described previously(7). After incubation at 37°C with 5% CO_2_ for 12-14 days, colony forming units (CFUs) were enumerated. To prevent analysis of extracellular replication, CFUs were not monitored after the onset of macrophage lysis.

### CBA and ELISA assays

BMDMs were seeded at 3X10^5^ cells/well in 48-well plates (TC-treated), and infected with *Hc* in triplicate (MOI=10 for CBA and MOI=2 for C3 ELISA). Supernatants were collected at the indicated times and either used fresh for assays or snap-frozen in LN_2_ and stored at −80°C. Supernatants were filtered using Spin-X cellulose acetate spin filters (Costar) by centrifugation. TNF-α was measured using the mouse TNF CBA flex set (BD) according to the manufacturer’s instructions. Data were acquired using a BD LSRII flow cytometer and analyzed using FCAP array software (BD). The colorimetric Mouse Complement C3 ELISA kit (Abcam) was used according to the manufacturer’s instructions to quantify C3 levels in macrophage culture supernatant. Mouse serum was incubated with *Hc* and zymosan at 10X10^8^ particles/mL for 30 min at 37°C. The reaction was stopped by addition of 10 mM EDTA and incubation on ice. *Hc* and zymosan were pelleted by centrifugation, and the supernatant was filtered using Spin-X cellulose acetate filters. Supernatants were diluted 1:200. A mouse C3a ELISA pair (BD) was used as previously described(97) according to manufacturer’s instructions to measure C3a levels in the supernatants. Corning High-Bind plates were coated with 4 μg/mL capture antibody in pH 6.5 binding buffer. PBS+10% FBS was used for blocking, and PBS+Δ10% FBS +0.05% Tween-20 was used to dilute samples, standards, and detection antibody solutions. Biotinylated C3a detection antibody was used at 6.25 ng/mL, and avidin-HRP was used at a 1:5000 dilution.

### Ethics statement

All mouse experiments were performed in compliance with the *National Institutes of Health Guide for the Care and Use of Laboratory Animals* and were approved by the Institutional Animal Care and Use Committee at the University of California San Francisco (protocol AN18753-03A). Mice were euthanized by CO_2_ narcosis and cervical dislocation consistent with American Veterinary Medical Association guidelines.

---

**Figure S1:**
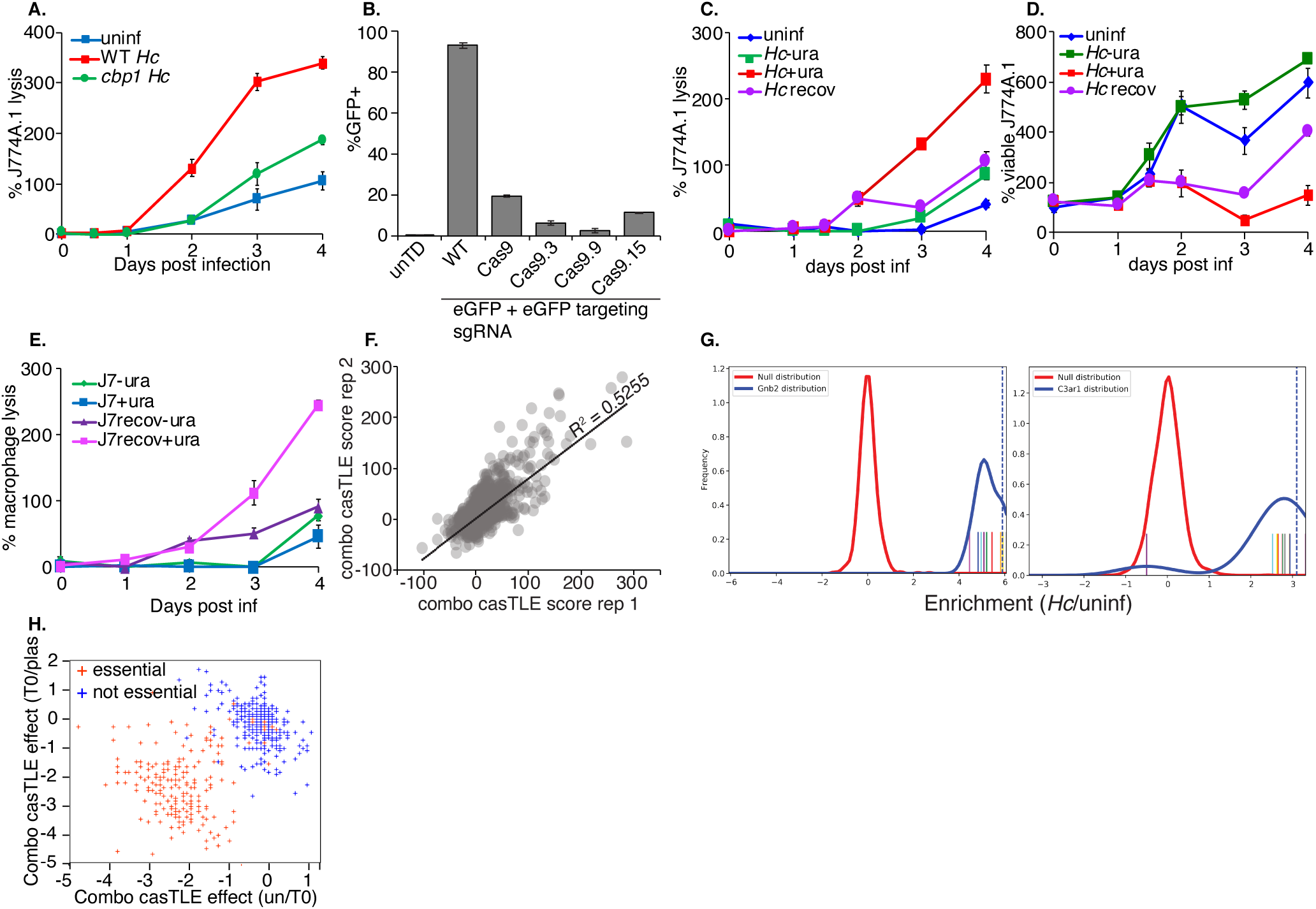
Development and validation of Cas9-expressing J7 cell-lines, and validation of screening approach. **A.** Characterization of *Hc-*mediated lysis in J774A.1 macrophage-like cells. J774A.1 cells were infected with WT *Hc,* or *Hc* with a disruption in a gene, *CBP1*, that is required for *Hc* to lyse macrophages. Lysis over time was measured using the LDH release assay. **B.** Validation and clonal expansion of Cas9-expressing J774A.1 cells. Cells were transduced with an Ef1a-Cas9-Blast expression vector and grown under blasticidin selection to generate a population of Cas9-expressing cells. These were subjected to single-cell sorting and clonal expansion to generate Cas9-expressing J774A.1 clones with high Cas9 activity. Cas9 activity was measured by transducing J774A.1 cells with a guide RNA vector that co-expressed EGFP with a sgRNA targeting EGFP. Cas9 activity leads to silencing of the GFP following puromycin selection. Cas9 clone 9 was chosen for the large-scale CRISPR screens due to its high-efficiency GFP silencing. **C-D.** Characterizing lysis and recovery from infection with uracil pulses during infection with a Ura5-deficient *Hc*. J774A.1 macrophages were infected with *ura5* mutant *Hc* in the presence or absence of exogenous uracil (0.4ug/mL). Uracil-containing cells were washed and media was replaced with uracil-poor media after 2d of lysis, which allowed the macrophages to recover. Recovery was assessed using LDH release quantification to assess lysis, and the confluency of viable cells in the wells was estimated using the pico-green dsDNA assay kit following lysis of macrophages with water. **E.** macrophages that had been recovered from lysis by removal of uracil from culture media were passaged for several days, and uracil was added to selected wells. Macrophage lysis over time was monitored by assessing LDH release over time to determine whether dormant yeast would be able to re-activate upon introduction of uracil. **F.** Reproducibility of the casTLE score across two replicates of the screens. **G.** Histograms comparing the distribution of negative control sgRNAs and sgRNAs targeting *Gnb2* or *C3ar* in the *H. capsulatum* infected pool compared to the uninfected pool. **H.** Analysis of essential gene behavior during J7 library growth. Scatter plot showing the gene effect resulting from passaging of J7s, either going from the plasmid pool to the T0 pool, or the T0 pool to the uninfected pool. Genes annotated as “essential” or “non-essential” were plotted to determine whether essential genes appeared more likely to drop out of the uninfected pools.

**Figure S2:**
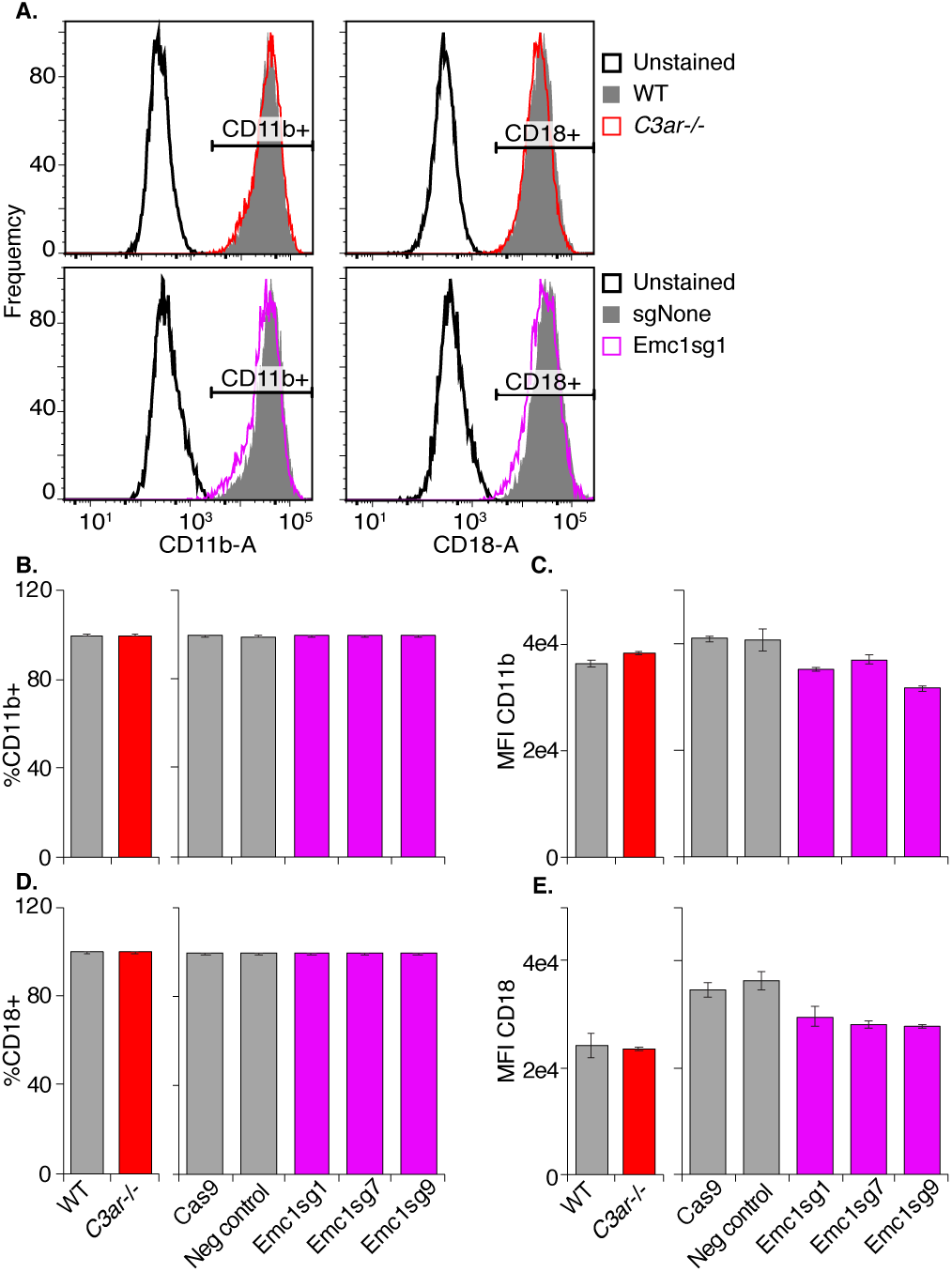
Emc1 and C3aR are not required for surface expression of CD18 or CD11b. BMDMs from *C3ar-/-* and WT mice, in addition to BMDMs expressing Cas9 and control or *Emc1-*targeting sgRNAs, were stained with anti-CD18 and anti-CD11b antibodies and analyzed by flow cytometry (n=2 biological replicates). **A.** Representative histograms showing CD11b and CD18 levels in control, *C3ar-/-*, and *Emc1* CRISPRKO BMDMs. The percentage of CD11b (**B**) and CD18 (**D**) positive macrophages was analyzed. The mean fluorescence intensity of CD11b (**C**) and CD18 (**D**) were also measured.

**Figure. S3:**
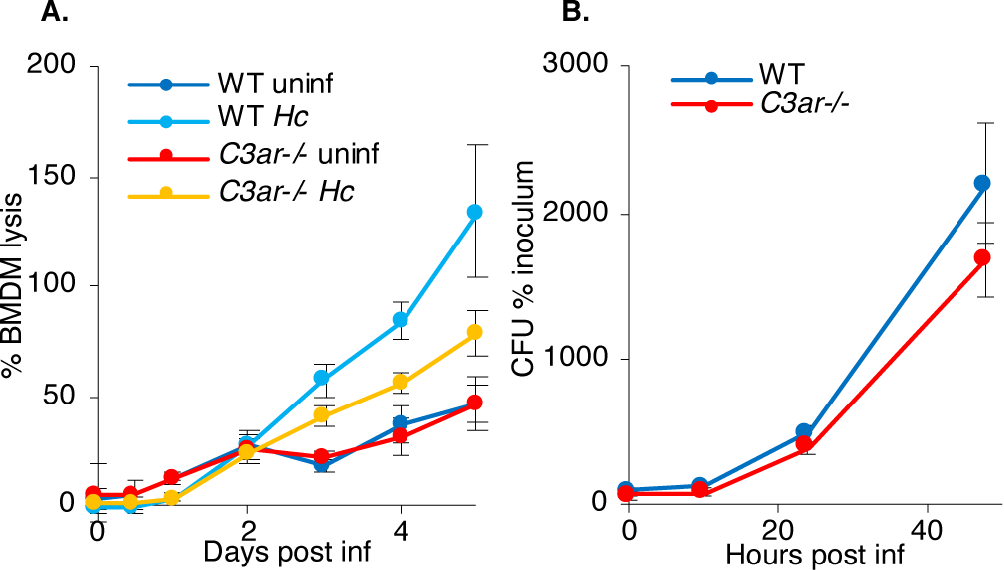
*C3ar-/-* BMDMs are partially resistant to *Hc-*induced lysis. BMDMs were infected with *Hc* (MOI=0.5), and macrophage lysis was quantified by measuring the release of lactate dehydrogenase into the culture supernatants over-time (n=3 biological replicates, 3 technical replicates/biorep) (**A**). At the indicated time points, macrophages were lysed using water, and lysates were spread on agar plates. Colony forming units (CFUs) were enumerated (n=3 biological replicates, 2 technical reps/biorep) (**B**).

**Figure. S4:**
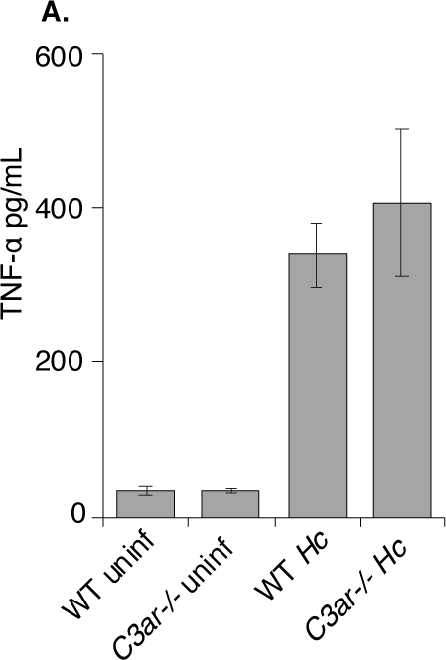
C3aR does not affect *Hc*-induced cytokine secretion by BMDMs. WT and *C3ar-/-* BMDMs were infected with *Hc* (MOI=5 for 6 h), and TNFα levels in macrophage supernatants were measured using the BD Cytometric Bead Array kit (n=3 biological replicates).

**Figure S5:**
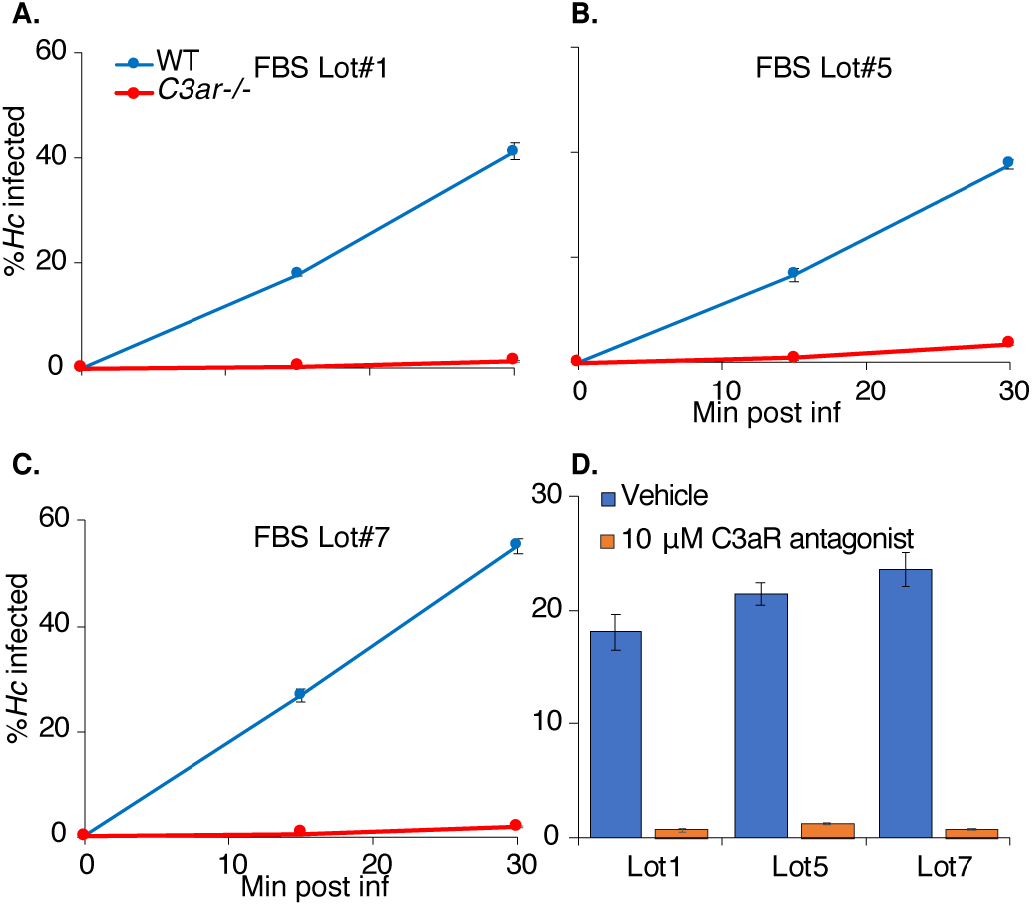
Different lots of FBS stimulate BMDM phagocytosis of *Hc* in a C3aR-dependent manner. (**A-C**) BMDMs from WT and *C3ar-/-* mice were infected with *Hc* in the presence of 20% FBS from three different lots from 2 separate suppliers. In addition, WT BMDMs differentiated in different lots of serum were treated with 10 µM of the C3aR antagonist and infected with *Hc* (**B**). Phagocytosis of *Hc* was measured by flow cytometry as described previously (n=2 biological replicates).

**Figure S6:**
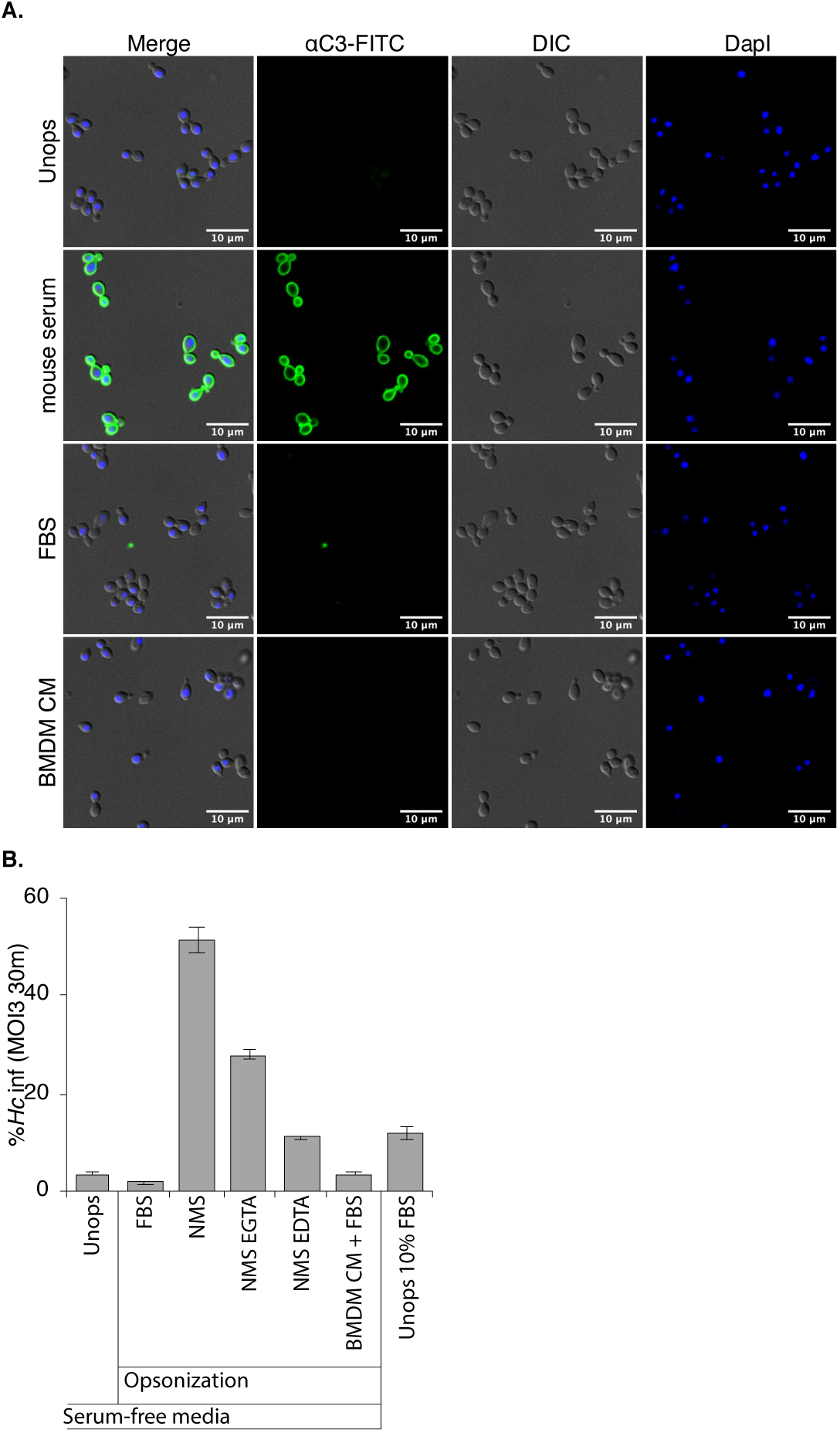
Macrophage conditioned media containing FBS does not promote opsonization that facilitates macrophage phagocytosis of *Hc* yeast in the absence of serum. BMDMs were cultured in media containing 10% FBS, and the BMDM conditioned media was harvested. *Hc* was incubated with macrophage conditioned media (BMDM CM), 10% FBS, or 10% normal mouse serum (NMS) with 10 mM EGTA or EDTA as indicated for 30 min 37°C. A. Incubation with conditioned media or FBS does not lead to C3 deposition on the *Hc* surface. C3 deposition on *Hc* yeast was analyzed by immunofluorescence microscopy using an anti-C3 antibody. B. Pre-incubation of yeast with conditioned media does not improve macrophage phagocytosis of *Hc.* Yeast were washed 2X and used to infect BMDMs at an MOI of 3 for 30 min in serum-free media. Phagocytosis was assessed using flow cytometry as previously described (n=3 biological replicates).

**Figure S7:**
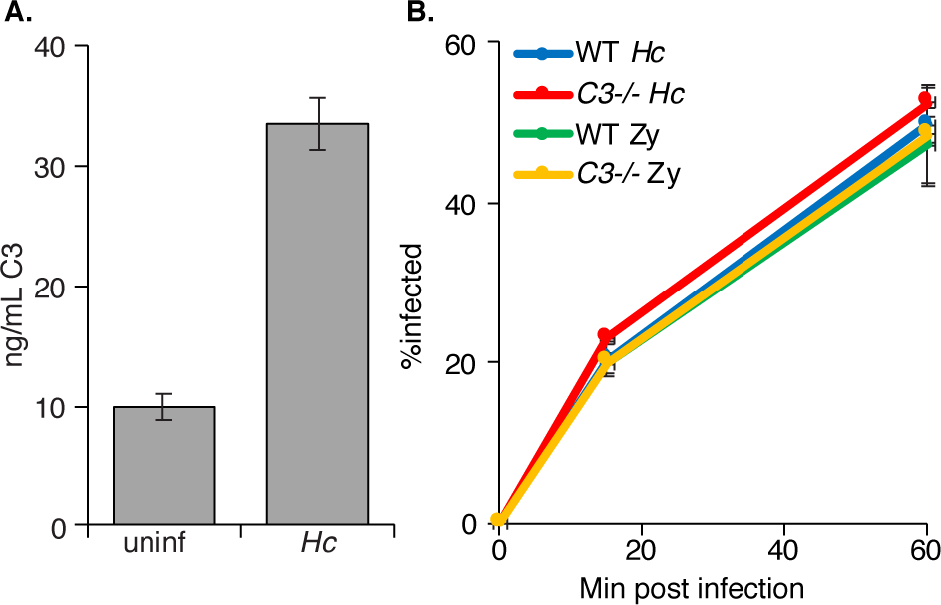
BMDM-derived C3 is not required for phagocytosis of *H. capsulatum* yeast or zymosan particles. **A.** BMDMs secrete C3 following infection with *Hc.* BMDMs were infected with *Hc* at an MOI2 for 24h, supernatants were harvested and C3 levels were quantified using a BD mouse C3 ELISA kit. **B.** *C3-/-* BMDMs are not defective in phagocytosis of *Hc* yeast or zymosan. WT and *C3-/-* BMDMs were infected with mCherry-expressing *H. capsulatum* or FITC-labelled zymosan, and uptake over time was measured using flow-cytometry. Extracellular yeasts were excluded with calcofluor white staining.

**Figure S8:**
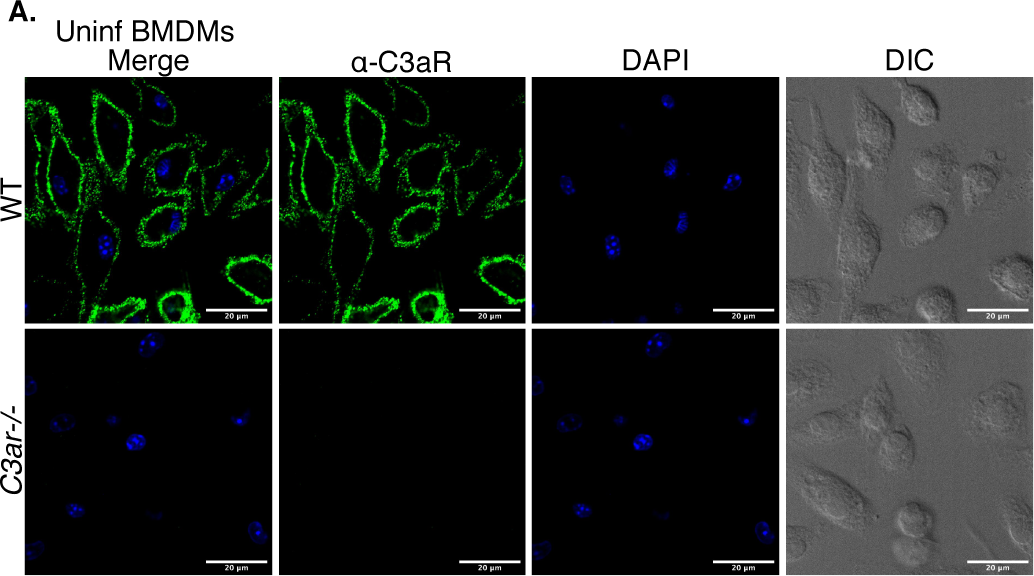
C3aR is localized to the plasma membrane in uninfected cells. Uninfected WT and *C3ar-/-* BMDMs were stained with a C3aR-specific antibody and imaged using confocal microscopy and optical sectioning. Representative slices of 2 biological replicates are shown. The antibody specifically detects C3aR, as staining was not observed in *C3ar-/-* BMDMs. C3aR exhibits punctate localization near the plasma membrane in WT BMDMs.

**Figure S9:**
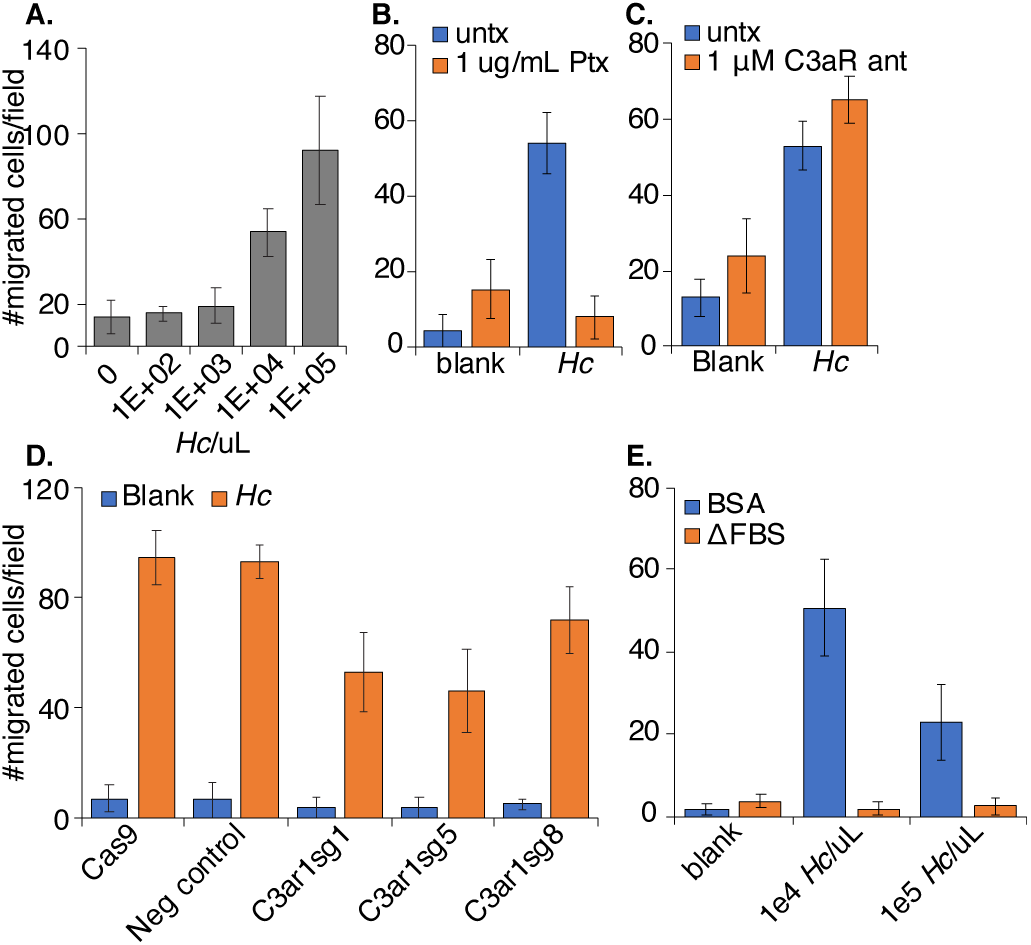
Macrophage-like cells undergo chemotaxis towards *H. capsulatum* yeast in a serum-independent manner, which is dependent on Gai, and partially dependent on C3aR. **A.** *Hc* stimulates chemotaxis of J774A.1 macrophage-like cells. *H. capsulatum* yeast were seeded into multiple-well plates at varying concentrations, and WT J774A.1 cells were seeded onto transwell permeable supports with 5 μm pores. Serum-free media supplemented with 0.25% BSA was used as the diluent in both the chamber and well unless otherwise indicated. After 3 h of migration, transwells were stained with crystal violet, and non-migratory cells were wiped off of the upper side of the transwell using a Q-tip. The number of migratory cells in each condition was quantified by microscopy (n=2 biogical replicates, 3 fields/biological replicate). B. Migration towards *Hc* is Gai-dependent. J774A.1 cells with or without pre-treatment with 1 µg/mL pertussis toxin (PTX) for 2 h were seeded into transwell permeable supports and migration towards 1e5 Hc/uL was quantified as described above. The number of migrating cells was quantified as described. C. The C3aR antagonist does not inhibit macrophage migration towards *Hc.* J774A.1 macrophages were treated with 1 µM SB290157, a C3aR antagonist, and migration towards *H. capsulatum* was assessed as described. D. C3aR-deficiency moderately impacts migration of J774A.1 cells towards *Hc.* Cas9-expressing J774A.1 macrophages transduced with non-targeting or C3aR-targeting sgRNAs were assessed for their ability to migrate towards *Hc* as described previously. E. *Hc-*dependent migration is abolished in the presence of FBS. The transwell migration assay was performed with media supplemented with BSA or 10% FBS to determine whether FBS affected the migration of macrophage-like cells towards *Hc* yeast.

**Figure S10:**
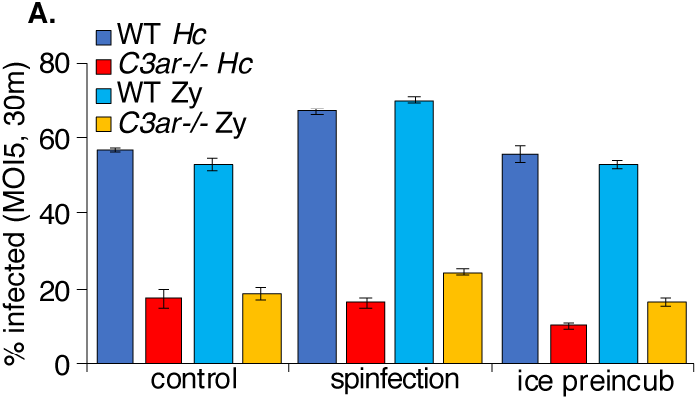
pre-incubation on ice or spinfection do not rescue phagocytosis of fungi in C3aR-/- BMDMs. **A.** BMDMs were infected with *Hc* or zymosan at an MOI=5 for 30 min. For the control condition, particles were added to the wells and allowed to settle onto the monolayer without intervention. For the 5 min spinfection, particles were added to the cells, and the plate was spun for 5 min at 550XG at RT before transferring to a 37°C, 5% CO_2_ incubator. For the ice preincubation condition, BMDMs were pre-chilled for 20 min on ice, and particles were allowed to settle onto the monolayer for 1 h on ice, then were transferred to a tissue culture incubator. Phagocytosis was measured as described previously (n=3 biological replicates).

**Figure S11:**
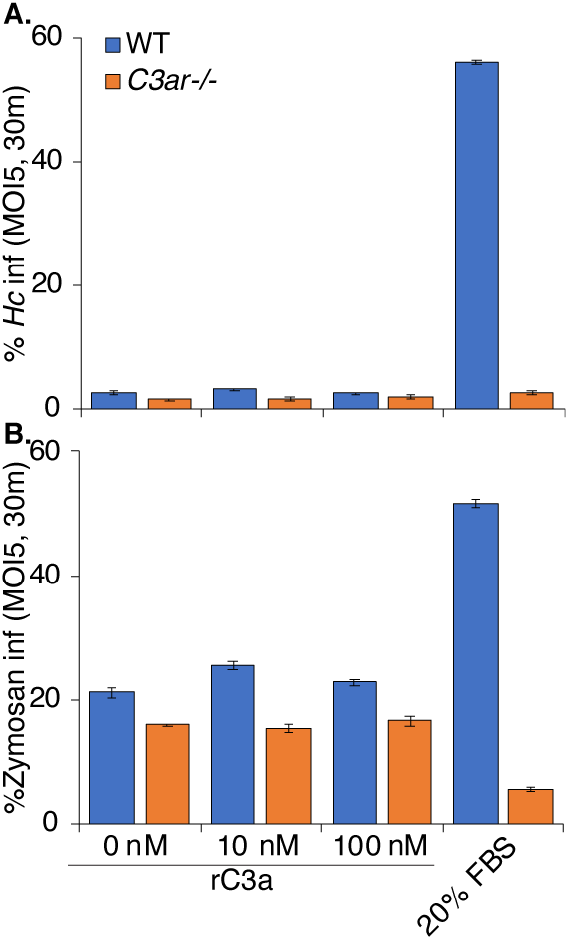
Addition of recombinant C3a is not sufficient to rescue phagocytosis of fungi in the absence of serum. BMDMs were pre-incubated in serum-free media with varying concentrations of recombinant mouse C3a (R&D systems) or with 20% FBS for 1 h, then infected with **A.** *Hc* or **B.** Zymosan at an MOI of 5 for 30 min. Phagocytosis was assessed by flow cytometry (n=2 biological replicates).

**Table S1:** results from CRISPR screens

Upload data (CSVs with results from castLE analysis, either separated or concatenated)

**Table S2:** Screen statistics summary by sub-library

| Sub-library | Rounds of *Hc*-mediated lysis | Duration (days) | Δdivisions (uninf-*Hc* inf) | Correlation coefficient ($R^2$ castLE score) | #genes | # desensitizing hits (*Hc*/un>0) | # sensitizing hits (*Hc*/un<0) | Total hits |
| --- | --- | --- | --- | --- | --- | --- | --- | --- |
| ACOCA | 2 | 16 | 10.17 | 0.74 | 1568 | 31 | 1 | 32 |
| ACOCB | 3 | 10 | 7.93 | 0.57 | 1546 | 59 | 1 | 60 |
| TMMA | 2 | 9 | 9.9 | 0.69 | 972 | 43 | 4 | 47 |
| TMMB | 3 | 7 | 6.73 | 0.49 | 951 | 37 | 1 | 38 |
| DTKPA | 2 | 7 | 5.53 | 0.24 | 1008 | 20 | 1 | 21 |
| DTKPB | 2 | 8 | 4.9 | 0.48 | 1003 | 15 | 0 | 15 |
| SPA | 2 | 9 | 8.5 | 0.49 | 1333 | 31 | 9 | 40 |
| SPB | 2 | 9 | 8.45 | 0.68 | 1478 | 21 | 2 | 23 |
| MPAB | 3 | 10 | 6.2 | 0.73 | 953 | 14 | 4 | 18 |
| GEA | 2 | 8 | 5.6 | 0.54 | 954 | 9 | 6 | 15 |
| GEB | 2 | 9 | 6.2 | 0.46 | 962 | 14 | 4 | 18 |
| U1A | 2 | 8 | 5.2 | 0.19 | 1019 | 8 | 1 | 9 |
| U1B | 2 | 8 | 5.3 | 0.53 | 932 | 19 | 2 | 21 |
| U2AB | 2 | 8 | 6.3 | 0.0028 | 2103 | 1 | 3 | 4 |

**Table S3:** sgRNAs used in this study

Upload CSV

**Table S4:** Oligonucleotides used in this study

Upload CSV

## Notes

### Competing Interest Statement

The authors have declared no competing interest.

## References

1. Pfaller MA, Diekema DJ. Epidemiology of invasive mycoses in North America. Critical reviews in microbiology. 2010;36(1):1–53.

2. Deepe GS, Jr., Gibbons RS, Smulian AG. Histoplasma capsulatum manifests preferential invasion of phagocytic subpopulations in murine lungs. J Leukoc Biol. 2008;84(3):669–78.

3. Jones GS, Sepúlveda VE, Goldman WE. Biodiverse Histoplasma Species Elicit Distinct Patterns of Pulmonary Inflammation following Sublethal Infection. mSphere. 2020;5(4).

4. Horwath MC, Fecher RA, Deepe GS, Jr. Histoplasma capsulatum, lung infection and immunity. Future microbiology. 2015;10(6):967–75.

5. Garfoot AL, Rappleye CA. Histoplasma capsulatum surmounts obstacles to intracellular pathogenesis. The FEBS journal. 2016;283(4):619–33.

6. Ray SC, Rappleye CA. Flying under the radar: Histoplasma capsulatum avoidance of innate immune recognition. Semin Cell Dev Biol. 2019;89:91–8.

7. English BC, Van Prooyen N, Ord T, Ord T, Sil A. The transcription factor CHOP, an effector of the integrated stress response, is required for host sensitivity to the fungal intracellular pathogen Histoplasma capsulatum. PLoS pathogens. 2017;13(9):e1006589.

8. Woods JP. Revisiting old friends: Developments in understanding Histoplasma capsulatum pathogenesis. Journal of Microbiology. 2016;54(3):265–76.

9. Isaac DT, Berkes CA, English BC, Hocking Murray D, Lee YN, Coady A, et al. Macrophage cell death and transcriptional response are actively triggered by the fungal virulence factor Cbp1 during H. capsulatum infection. Mol Microbiol. 2015;98(5):910–29.

10. Sebghati TS, Engle JT, Goldman WE. Intracellular parasitism by Histoplasma capsulatum: fungal virulence and calcium dependence. Science (New York, NY). 2000;290(5495):1368–72.

11. Erwig LP, Gow NA. Interactions of fungal pathogens with phagocytes. Nature reviews Microbiology. 2016;14(3):163–76.

12. Brown GD, Taylor PR, Reid DM, Willment JA, Williams DL, Martinez-Pomares L, et al. Dectin-1 is a major beta-glucan receptor on macrophages. The Journal of experimental medicine. 2002;196(3):407–12.

13. Goodridge HS, Reyes CN, Becker CA, Katsumoto TR, Ma J, Wolf AJ, et al. Activation of the innate immune receptor Dectin-1 upon formation of a ‘phagocytic synapse’. Nature. 2011;472(7344):471–5.

14. Flannagan RS, Jaumouillé V, Grinstein S. The cell biology of phagocytosis. Annu Rev Pathol. 2012;7:61–98.

15. Freeman SA, Grinstein S. Phagocytosis: receptors, signal integration, and the cytoskeleton. Immunological reviews. 2014;262(1):193–215.

16. Lin JS, Huang JH, Hung LY, Wu SY, Wu-Hsieh BA. Distinct roles of complement receptor 3, Dectin-1, and sialic acids in murine macrophage interaction with Histoplasma yeast. J Leukoc Biol. 2010;88(1):95–106.

17. Rappleye CA, Eissenberg LG, Goldman WE. Histoplasma capsulatum alpha-(1,3)-glucan blocks innate immune recognition by the beta-glucan receptor. Proceedings of the National Academy of Sciences of the United States of America. 2007;104(4):1366–70.

18. Garfoot AL, Shen Q, Wuthrich M, Klein BS, Rappleye CA. The Eng1 beta-Glucanase Enhances Histoplasma Virulence by Reducing beta-Glucan Exposure. mBio. 2016;7(2).

19. Bullock WE, Wright SD. ROLE OF THE ADHERENCE-PROMOTING RECEPTORS, CR3, LFA-1, AND P150,95, IN BINDING OF HISTOPLASMA-CAPSULATUM BY HUMAN MACROPHAGES. Journal of Experimental Medicine. 1987;165(1):195–210.

20. Tan SM. The leucocyte β2 (CD18) integrins: the structure, functional regulation and signalling properties. Biosci Rep. 2012;32(3):241–69.

21. Noris M, Remuzzi G. Overview of complement activation and regulation. Semin Nephrol. 2013;33(6):479–92.

22. Wong SSW, Daniel I, Gangneux JP, Jayapal JM, Guegan H, Dellière S, et al. Differential Interactions of Serum and Bronchoalveolar Lavage Fluid Complement Proteins with Conidia of Airborne Fungal Pathogen Aspergillus fumigatus. Infect Immun. 2020;88(9).

23. Bolger MS, Ross DS, Jiang H, Frank MM, Ghio AJ, Schwartz DA, et al. Complement levels and activity in the normal and LPS-injured lung. Am J Physiol Lung Cell Mol Physiol. 2007;292(3):L748–59.

24. Lambris JD, Ricklin D, Geisbrecht BV. Complement evasion by human pathogens. Nature reviews Microbiology. 2008;6(2):132–42.

25. Torres-Gomez A, Cabañas C, Lafuente EM. Phagocytic Integrins: Activation and Signaling. Front Immunol. 2020;11:738.

26. Quell KM, Karsten CM, Kordowski A, Almeida LN, Briukhovetska D, Wiese AV, et al. Monitoring C3aR Expression Using a Floxed tdTomato-C3aR Reporter Knock-in Mouse. Journal of immunology (Baltimore, Md : 1950). 2017;199(2):688–706.

27. Zwirner J, Werfel T, Wilken HC, Theile E, Gotze O. Anaphylatoxin C3a but not C3a(desArg) is a chemotaxin for the mouse macrophage cell line J774. European journal of immunology. 1998;28(5):1570–7.

28. Coulthard LG, Woodruff TM. Is the complement activation product C3a a proinflammatory molecule? Re-evaluating the evidence and the myth. Journal of immunology (Baltimore, Md : 1950). 2015;194(8):3542–8.

29. Klos A, Tenner AJ, Johswich KO, Ager RR, Reis ES, Köhl J. The role of the anaphylatoxins in health and disease. Mol Immunol. 2009;46(14):2753–66.

30. Ezekowitz RA, Sim RB, Hill M, Gordon S. Local opsonization by secreted macrophage complement components. Role of receptors for complement in uptake of zymosan. The Journal of experimental medicine. 1984;159(1):244–60.

31. Goodrum KJ. Complement component C3 secretion by mouse macrophage-like cell lines. J Leukoc Biol. 1987;41(4):295–301.

32. Luo C, Chen M, Madden A, Xu H. Expression of complement components and regulators by different subtypes of bone marrow-derived macrophages. Inflammation. 2012;35(4):1448–61.

33. Galgiani JN, Yam P, Petz LD, Williams PL, Stevens DA. Complement activation by Coccidioides immitis: in vitro and clinical studies. Infect Immun. 1980;28(3):944–9.

34. Ratnoff WD, Pepple JM, Winkelstein JA. Activation of the alternative complement pathway by Histoplasma capsulatum. Infect Immun. 1980;30(1):147–9.

35. Kozel TR. Activation of the complement system by pathogenic fungi. Clinical microbiology reviews. 1996;9(1):34–46.

36. Zaragoza O, Taborda CP, Casadevall A. The efficacy of complement-mediated phagocytosis of Cryptococcus neoformans is dependent on the location of C3 in the polysaccharide capsule and involves both direct and indirect C3-mediated interactions. European journal of immunology. 2003;33(7):1957–67.

37. Harpf V, Rambach G, Würzner R, Lass-Flörl C, Speth C. Candida and Complement: New Aspects in an Old Battle. Front Immunol. 2020;11:1471.

38. Taylor PR, Tsoni SV, Willment JA, Dennehy KM, Rosas M, Findon H, et al. Dectin-1 is required for beta-glucan recognition and control of fungal infection. Nature immunology. 2007;8(1):31–8.

39. Tsoni SV, Kerrigan AM, Marakalala MJ, Srinivasan N, Duffield M, Taylor PR, et al. Complement C3 plays an essential role in the control of opportunistic fungal infections. Infect Immun. 2009;77(9):3679–85.

40. Shapiro S, Beenhouwer DO, Feldmesser M, Taborda C, Carroll MC, Casadevall A, et al. Immunoglobulin G monoclonal antibodies to Cryptococcus neoformans protect mice deficient in complement component C3. Infect Immun. 2002;70(5):2598–604.

41. Pillemer L, Blum L, Lepow IH, Ross OA, Todd EW, Wardlaw AC. The properdin system and immunity. I. Demonstration and isolation of a new serum protein, properdin, and its role in immune phenomena. Science (New York, NY). 1954;120(3112):279–85.

42. Sun D, Zhang M, Liu G, Wu H, Zhu X, Zhou H, et al. Real-Time Imaging of Interactions of Neutrophils with Cryptococcus neoformans Demonstrates a Crucial Role of Complement C5a-C5aR Signaling. Infect Immun. 2016;84(1):216–29.

43. Cheng SC, Sprong T, Joosten LA, van der Meer JW, Kullberg BJ, Hube B, et al. Complement plays a central role in Candida albicans-induced cytokine production by human PBMCs. European journal of immunology. 2012;42(4):993–1004.

44. Kampmann M, Bassik MC, Weissman JS. Functional genomics platform for pooled screening and generation of mammalian genetic interaction maps. Nature Protocols. 2014;9(8):1825–47.

45. Jeng EE, Bhadkamkar V, Ibe NU, Gause H, Jiang L, Chan J, et al. Systematic Identification of Host Cell Regulators of Legionella pneumophila Pathogenesis Using a Genome-wide CRISPR Screen. Cell Host Microbe. 2019;26(4):551–63.e6.

46. Park RJ, Wang T, Koundakjian D, Hultquist JF, Lamothe-Molina P, Monel B, et al. A genome-wide CRISPR screen identifies a restricted set of HIV host dependency factors. Nat Genet. 2017;49(2):193–203.

47. Napier BA, Brubaker SW, Sweeney TE, Monette P, Rothmeier GH, Gertsvolf NA, et al. Complement pathway amplifies caspase-11-dependent cell death and endotoxin-induced sepsis severity. The Journal of experimental medicine. 2016;213(11):2365–82.

48. Morgens DW, Wainberg M, Boyle EA, Ursu O, Araya CL, Tsui CK, et al. Genome-scale measurement of off-target activity using Cas9 toxicity in high-throughput screens. Nature communications. 2017;8:15178.

49. Worsham PL, Goldman WE. Selection and characterization of ura5 mutants of Histoplasma capsulatum. Molecular & general genetics : MGG. 1988;214(2):348–52.

50. Morgens DW, Deans RM, Li A, Bassik MC. Systematic comparison of CRISPR/Cas9 and RNAi screens for essential genes. Nature biotechnology. 2016;34(6):634–6.

51. Haney MS, Bohlen CJ, Morgens DW, Ousey JA, Barkal AA, Tsui CK, et al. Identification of phagocytosis regulators using magnetic genome-wide CRISPR screens. Nat Genet. 2018;50(12):1716–27.

52. Devreotes P, Horwitz AR. Signaling networks that regulate cell migration. Cold Spring Harb Perspect Biol. 2015;7(8):a005959.

53. Moser M, Nieswandt B, Ussar S, Pozgajova M, Fässler R. Kindlin-3 is essential for integrin activation and platelet aggregation. Nat Med. 2008;14(3):325–30.

54. Jewell JL, Russell RC, Guan KL. Amino acid signalling upstream of mTOR. Nature reviews Molecular cell biology. 2013;14(3):133–9.

55. Chitwood PJ, Hegde RS. The Role of EMC during Membrane Protein Biogenesis. Trends Cell Biol. 2019.

56. Shurtleff MJ, Itzhak DN, Hussmann JA, Schirle Oakdale NT, Costa EA, Jonikas M, et al. The ER membrane protein complex interacts cotranslationally to enable biogenesis of multipass membrane proteins. Elife. 2018;7.

57. Irannejad R, von Zastrow M. GPCR signaling along the endocytic pathway. Current opinion in cell biology. 2014;27:109–16.

58. Block H, Stadtmann A, Riad D, Rossaint J, Sohlbach C, Germena G, et al. Gnb isoforms control a signaling pathway comprising Rac1, Plcβ2, and Plcβ3 leading to LFA-1 activation and neutrophil arrest in vivo. Blood. 2016;127(3):314–24.

59. Andreu N, Phelan J, de Sessions PF, Cliff JM, Clark TG, Hibberd ML. Primary macrophages and J774 cells respond differently to infection with Mycobacterium tuberculosis. Scientific reports. 2017;7:42225.

60. Haas PJ, van Strijp J. Anaphylatoxins: their role in bacterial infection and inflammation. Immunologic research. 2007;37(3):161–75.

61. Humbles AA, Lu B, Nilsson CA, Lilly C, Israel E, Fujiwara Y, et al. A role for the C3a anaphylatoxin receptor in the effector phase of asthma. Nature. 2000;406(6799):998–1001.

62. Mueller-Ortiz SL, Hollmann TJ, Haviland DL, Wetsel RA. Ablation of the complement C3a anaphylatoxin receptor causes enhanced killing of Pseudomonas aeruginosa in a mouse model of pneumonia. Am J Physiol Lung Cell Mol Physiol. 2006;291(2):L157–65.

63. Wu KY, Zhang T, Zhao GX, Ma N, Zhao SJ, Wang N, et al. The C3a/C3aR axis mediates anti-inflammatory activity and protects against uropathogenic E coli-induced kidney injury in mice. Kidney Int. 2019;96(3):612–27.

64. Lian H, Litvinchuk A, Chiang AC, Aithmitti N, Jankowsky JL, Zheng H. Astrocyte-Microglia Cross Talk through Complement Activation Modulates Amyloid Pathology in Mouse Models of Alzheimer’s Disease. The Journal of neuroscience : the official journal of the Society for Neuroscience. 2016;36(2):577–89.

65. Litvinchuk A, Wan YW, Swartzlander DB, Chen F, Cole A, Propson NE, et al. Complement C3aR Inactivation Attenuates Tau Pathology and Reverses an Immune Network Deregulated in Tauopathy Models and Alzheimer’s Disease. Neuron. 2018;100(6):1337–53.e5.

66. Zhang LY, Pan J, Mamtilahun M, Zhu Y, Wang L, Venkatesh A, et al. Microglia exacerbate white matter injury via complement C3/C3aR pathway after hypoperfusion. Theranostics. 2020;10(1):74–90.

67. Ames RS, Lee D, Foley JJ, Jurewicz AJ, Tornetta MA, Bautsch W, et al. Identification of a selective nonpeptide antagonist of the anaphylatoxin C3a receptor that demonstrates antiinflammatory activity in animal models. Journal of immunology (Baltimore, Md : 1950). 2001;166(10):6341–8.

68. Huang NN, Becker S, Boularan C, Kamenyeva O, Vural A, Hwang IY, et al. Canonical and noncanonical g-protein signaling helps coordinate actin dynamics to promote macrophage phagocytosis of zymosan. Molecular and cellular biology. 2014;34(22):4186–99.

69. Decker T, Lohmann-Matthes ML. A quick and simple method for the quantitation of lactate dehydrogenase release in measurements of cellular cytotoxicity and tumor necrosis factor (TNF) activity. J Immunol Methods. 1988;115(1):61–9.

70. Nilsson UR, Müller-Eberhard HJ. Deficiency of the fifth component of complement in mice with an inherited complement defect. The Journal of experimental medicine. 1967;125(1):1–16.

71. Zhang MX, Klein B. Activation, binding, and processing of complement component 3 (C3) by Blastomyces dermatitidis. Infect Immun. 1997;65(5):1849–55.

72. Riedl J, Crevenna AH, Kessenbrock K, Yu JH, Neukirchen D, Bista M, et al. Lifeact: a versatile marker to visualize F-actin. Nature methods. 2008;5(7):605–7.

73. Svensson CM, Medyukhina A, Belyaev I, Al-Zaben N, Figge MT. Untangling cell tracks: Quantifying cell migration by time lapse image data analysis. Cytometry Part A : the journal of the International Society for Analytical Cytology. 2018;93(3):357–70.

74. Rougerie P, Miskolci V, Cox D. Generation of membrane structures during phagocytosis and chemotaxis of macrophages: role and regulation of the actin cytoskeleton. Immunological reviews. 2013;256(1):222–39.

75. Van Haastert PJ, Devreotes PN. Chemotaxis: signalling the way forward. Nature reviews Molecular cell biology. 2004;5(8):626–34.

76. Peracino B, Borleis J, Jin T, Westphal M, Schwartz JM, Wu L, et al. G protein beta subunit-null mutants are impaired in phagocytosis and chemotaxis due to inappropriate regulation of the actin cytoskeleton. The Journal of cell biology. 1998;141(7):1529–37.

77. Pan M, Neilson MP, Grunfeld AM, Cruz P, Wen X, Insall RH, et al. A G-protein-coupled chemoattractant receptor recognizes lipopolysaccharide for bacterial phagocytosis. PLoS Biol. 2018;16(5):e2005754.

78. Freeman SA, Vega A, Riedl M, Collins RF, Ostrowski PP, Woods EC, et al. Transmembrane Pickets Connect Cyto- and Pericellular Skeletons Forming Barriers to Receptor Engagement. Cell. 2018;172(1-2):305–17.e10.

79. Flannagan RS, Harrison RE, Yip CM, Jaqaman K, Grinstein S. Dynamic macrophage “probing” is required for the efficient capture of phagocytic targets. The Journal of cell biology. 2010;191(6):1205–18.

80. Shen K, Sidik H, Talbot WS. The Rag-Ragulator Complex Regulates Lysosome Function and Phagocytic Flux in Microglia. Cell Rep. 2016;14(3):547–59.

81. Gudjonsson T, Altmeyer M, Savic V, Toledo L, Dinant C, Grøfte M, et al. TRIP12 and UBR5 suppress spreading of chromatin ubiquitylation at damaged chromosomes. Cell. 2012;150(4):697–709.

82. Cho JH, Kim SA, Seo YS, Park SG, Park BC, Kim JH, et al. The p90 ribosomal S6 kinase-UBR5 pathway controls Toll-like receptor signaling via miRNA-induced translational inhibition of tumor necrosis factor receptor-associated factor 3. The Journal of biological chemistry. 2017;292(28):11804–14.

83. Surugiu R, Catalin B, Dumbrava D, Gresita A, Olaru DG, Hermann DM, et al. Intracortical Administration of the Complement C3 Receptor Antagonist Trifluoroacetate Modulates Microglia Reaction after Brain Injury. Neural Plast. 2019;2019:1071036.

84. Sun D, Sun P, Li H, Zhang M, Liu G, Strickland AB, et al. Fungal dissemination is limited by liver macrophage filtration of the blood. Nature communications. 2019;10(1):4566.

85. Hwang LH, Mayfield JA, Rine J, Sil A. Histoplasma requires SID1, a member of an iron-regulated siderophore gene cluster, for host colonization. PLoS pathogens. 2008;4(4):e1000044.

86. Van Prooyen N, Henderson CA, Hocking Murray D, Sil A. CD103+ Conventional Dendritic Cells Are Critical for TLR7/9-Dependent Host Defense against Histoplasma capsulatum, an Endemic Fungal Pathogen of Humans. PLoS pathogens. 2016;12(7):e1005749.

87. Worsham PL, Goldman WE. Quantitative plating of Histoplasma capsulatum without addition of conditioned medium or siderophores. Journal of medical and veterinary mycology : bi-monthly publication of the International Society for Human and Animal Mycology. 1988;26(3):137–43.

88. Mead HL, Van Dyke MCC, Barker BM. Proper Care and Feeding of Coccidioides: A Laboratorian’s Guide to Cultivating the Dimorphic Stages of C. immitis and C. posadasii. Curr Protoc Microbiol. 2020;58(1):e113.

89. Deans RM, Morgens DW, Okesli A, Pillay S, Horlbeck MA, Kampmann M, et al. Parallel shRNA and CRISPR-Cas9 screens enable antiviral drug target identification. Nat Chem Biol. 2016;12(5):361–6.

90. Brinkman EK, Chen T, Amendola M, van Steensel B. Easy quantitative assessment of genome editing by sequence trace decomposition. Nucleic acids research. 2014;42(22):e168.

91. Bassik MC, Kampmann M, Lebbink RJ, Wang SY, Hein MY, Poser I, et al. A Systematic Mammalian Genetic Interaction Map Reveals Pathways Underlying Ricin Susceptibility. Cell. 2013;152(4):909–22.

92. Saldanha AJ. Java Treeview--extensible visualization of microarray data. Bioinformatics. 2004;20(17):3246–8.

93. Eden E, Navon R, Steinfeld I, Lipson D, Yakhini Z. GOrilla: a tool for discovery and visualization of enriched GO terms in ranked gene lists. BMC bioinformatics. 2009;10:48.

94. Ollion J, Cochennec J, Loll F, Escudé C, Boudier T. TANGO: a generic tool for high-throughput 3D image analysis for studying nuclear organization. Bioinformatics. 2013;29(14):1840–1.

95. Meijering E, Dzyubachyk O, Smal I. Methods for cell and particle tracking. Methods Enzymol. 2012;504:183–200.

96. Isaac DT, Coady A, Van Prooyen N, Sil A. The 3-Hydroxy-Methylglutaryl Coenzyme A Lyase HCL1 Is Required for Macrophage Colonization by Human Fungal Pathogen Histoplasma capsulatum. Infection and Immunity. 2013;81(2):411–20.

97. Roy RM, Paes HC, Nanjappa SG, Sorkness R, Gasper D, Sterkel A, et al. Complement component 3C3 and C3a receptor are required in chitin-dependent allergic sensitization to Aspergillus fumigatus but dispensable in chitin-induced innate allergic inflammation. mBio. 2013;4(2).

